# Distinct mechanisms of non-autonomous UPR^ER^ mediated by GABAergic, glutamatergic, and octopaminergic neurons

**DOI:** 10.1101/2024.05.27.595950

**Authors:** Aeowynn J. Coakley, Adam J. Hruby, Jing Wang, Andrew Bong, Tripti Nair, Carmen M. Ramos, Athena Alcala, Daniel Hicks, Maxim Averbukh, Naibedya Dutta, Darius Moaddeli, Cynthia Siebrand, Mattias de los Rios Rogers, Arushi Sahay, Sean P. Curran, Peter J. Mullen, Bérénice A Benayoun, Gilberto Garcia, Ryo Higuchi-Sanabria

## Abstract

The capacity to deal with stress declines during the aging process, and preservation of cellular stress responses is critical to healthy aging. The unfolded protein response of the endoplasmic reticulum (UPR^ER^) is one such conserved mechanism, which is critical for the maintenance of several major functions of the ER during stress, including protein folding and lipid metabolism. Hyperactivation of the UPR^ER^ by overexpression of the major transcription factor, *xbp-1s*, solely in neurons drives lifespan extension as neurons send a neurotransmitter-based signal to other tissue to activate UPR^ER^ in a non-autonomous fashion. Previous work identified serotonergic, dopaminergic, and tyraminergic neurons in this signaling paradigm. To further expand our understanding of the neural circuitry that underlies the non-autonomous signaling of ER stress, we activated UPR^ER^ solely in glutamatergic, octopaminergic, and GABAergic neurons in *C. elegans* and paired whole-body transcriptomic analysis with functional assays. We found that UPR^ER^-induced signals from glutamatergic neurons increased expression of canonical protein homeostasis pathways and octopaminergic neurons promoted pathogen response pathways; while minor, statistically significant changes were observed in lipid metabolism-related genes with GABAergic UPR^ER^ activation. These findings provide further evidence for the distinct role neuronal subtypes play in driving the diverse response to ER stress.

## Introduction

Regulation of organelle homeostasis is essential for maintenance of cellular health, which has direct implications for organismal health and longevity. The endoplasmic reticulum (ER) is one such organelle, which processes about a third of the proteins and lipids in cells and has dedicated quality control machineries to preserve these and numerous other functions. One primary quality control machinery is the ER unfolded protein response (UPR^ER^), a transcriptional response to ER damage or stress, which activates genes essential for maintenance of proper ER function^1^. Activation of the UPR^ER^ involves the ER-membrane protein, inositol-requiring enzyme 1 (IRE-1), which dimerizes upon sensing ER stress to splice X-box binding protein (*xbp-1*) mRNA into *xbp-1s*. *xbp-1s* mRNA encodes a functional transcription factor, XBP-1s, which activates genes essential for restoring ER homeostasis, including protein chaperones, autophagy, and the ubiquitin proteosome system, among others. This transcriptional response to stress is essential for maintaining proper function of the ER and has direct implications in longevity. Specifically, UPR^ER^ function has been shown to decline with age, and heightened activation of UPR^ER^ maintained stress resilience at old age in *C. elegans*^2^.

Overexpression of *xbp-1s* solely in neurons is sufficient to enhance *C. elegans* lifespan due to a whole-body UPR^ER^ activation through neuron-to-body communication mediated by neurotransmitter signaling^2^. Upon neuronal UPR^ER^ activation, a complex signaling event mediated by a combination of dopamine, serotonin^3^, and tyramine^4^, results in dramatic remodeling of peripheral cells. Specifically, intestinal cells can activate proteostasis^4^, lipid metabolism^5,6^, and lysosomal function^7^ to drive longevity. These studies revealed numerous neuronal subtypes and distinct mechanistic pathways, including chaperone induction downstream of serotonergic signaling, lipid remodeling through lipophagy downstream of dopaminergic signaling, and proteostasis machinery through tyramine signaling from RIM and RIC neurons, all of which are essential to promote longevity. Finally, this UPR^ER^ signaling is not limited to neurons, as several glial subtypes were also capable of eliciting a glia-to-body UPR^ER^ signaling event to promote longevity^8^.

Similar homeostatic benefits of UPR^ER^ in neurons are observed in mammals, wherein *Xbp1s* expression in pro-opiomelanocortin (POMC) neurons has been shown to protect against diet-induced obesity by improving leptin and insulin sensitivity under ER stress^9^. While all these studies utilized an artificial transgenic expression system, two recent studies have shown that neuron-to-body UPR^ER^ signaling is also occurs in endogenous pathways. In mice, olfactory perception of food is sufficient to promote POMC *Xbp1s* expression and activation of post- prandial liver ER adaption^10^. In *C. elegans*, chemosensation of pathogenic bacteria was found to promote neuronal *xbp1-s* expression, leading to UPR^ER^ activation in peripheral tissues and extension of lifespan^11^. These studies revealed that endogenous neuron-to-body signaling utilized similar mechanistic pathways to *xbp-1s* overexpression paradigms, which highlight the translatability of using transgenic approaches to dissect the neuronal circuitry of UPR^ER^ signaling.

Building on previous research, we were interested in understanding whether other neuronal subtypes are involved in neuron-to-body UPR^ER^ activity. We sought to determine whether glutamatergic, GABAergic, and octopaminergic neurons are necessary and/or sufficient to drive neuron-to-body communication of the UPR^ER^ in *C. elegans*. We accomplished this by *xbp-1s* overexpression in these neuronal subtypes and assessing measurements of general health, such as lifespan, healthspan, ER function, and stress resilience. Further, we performed a comprehensive transcriptomic analysis to identify potential mechanistic pathways that drive phenotypic outcomes in these neuronal subtype UPR^ER^ paradigms.

## Materials and Methods

### *C. elegans* maintenance

All strains utilized in this investigation are derived from the N2 wild-type worm sourced from the Caenorhabditis Genetics Center (CGC) and are detailed below. The worms are maintained at 15°C, fed with OP50 *E. coli* B strain. Animals are bleached and L1 arrested as described below for all experimentation and transferred to growth conditions at 20°C, utilizing HT115 *E. coli* K strain for all experiments. Experiments employed HT115 bacteria carrying an empty pL4440 vector referred to as empty vector (EV). An important note is that the *tbh-1p::xbp-1s* animals represent a bimodal population, where some animals have severely stunted growth. Those with stunted growth also exhibit a significant decrease in lifespan, so for the majority of the manuscript, we opted to select for the animals that are similar in size to wild-type animals, which we call “normal” sized for ease. Wherever relevant, data is represented as “mixed” when we did not separate the “normal” sized versus the “small” sized animals and labeled as “small” when animals with stunted growth were isolated. Wherever data is presented simply as “*tbh-1*”, these are the “normal” sized animals.

### Plates

Standard NGM plates for maintenance using OP50 contained the following: Bacto-Agar (Difco) 2% w/v, Bacto Peptone 0.25% w/v, NaCl_2_ 0.3% w/v, 1 mM CaCl_2_, 5 µg/ml cholesterol, 0.625 mM KPO_4_ pH 6.0, 1 mM MgSO_4_.

Standard NGM plates for experimental condition using HT115 bacteria contained the following: 2% RNAi plates for experiments contained the following: Bacto-Agar (Difco) 2% w/v, Bacto Peptone 0.25% w/v, NaCl_2_ 0.3% w/v, 1 mM CaCl_2_, 5 µg/ml cholesterol, 0.625 mM KPO_4_ pH 6.0, 1 mM MgSO_4_, 100 µg/mL carbenicillin, 1 mM IPTG.

For all aging experiments, 100 μL of 10 mg/mL (+)-5-Fluorodeoxyuridine (FUDR) was placed directly on the bacterial lawn and animals are moved onto FUDR-containing plates at day 1 of adulthood. FUDR is not used for plates containing tunicamycin, as tunicamycin prohibits growth of progeny and thus use of FUDR is not necessary^12^.

### Bleaching

Experiments were conducted on animals of the same age, synchronized using a standard bleaching protocol. Worms were collected into a 15 mL conical tube using M9 solution (22 mM KH_2_PO_4_ monobasic, 42.3 mM NaHPO_4_, 85.6 mM NaCl, 1 mM MgSO_4_) and subjected to a bleaching solution (1.8% sodium hypochlorite, 0.375 M NaOH in M9) until complete digestion of carcasses. Intact eggs were then washed four times with M9 solution by centrifugation at 1,100 x g for 30 seconds. After the final wash, animals were L1 arrested by incubating overnight in M9 at 20°C on a rotator for a maximum of 24 hours.

### Transgenic strain synthesis

The sequence for *xbp-1s* expression was defined as previously described^2^ and is provided below. The *xbp-1s* coding sequence was cloned from cDNA synthesized via reverse transcriptase using RNA isolated from N2 worms, the endogenous *eat-4*, *tbh-1*, and *unc-25* promoter was cloned from gDNA isolated from N2 worms, and an *unc-54* 3′UTR was cloned from gDNA isolated from N2 worms. Plasmids were injected into N2 worms using a standard microinjection protocol as described^13^ with 10 ng/μl of overexpression plasmid, 2.5 ng/μl of *myo- 2p::mCherry* or *myo-2p::GFP* as a co-injection marker. Both injections and integration of constructs were performed by SUNY Biotech. All integrated animals were then backcrossed to our N2 lines >8 times to eliminate mutations and create an isogenic line. All sequences used in this manuscript are as follows:

*xbp-1s* ATGAGCAACTATCCAAAACGTATTTATGTGCTCCCAGCACGCCACGTGGCAGCGCCACAGCCTCAGAGAATGGCTCCCAAGCGTGCACTTCCAACAGAACAAGTTGTCGCACAACTTCTTGGCGATGATATGGGACCATCTGGGCCACGCAAAAGAGAACGACTGAATCATTTGAGTCAGGAGGAGAAAATGGATCGTCGGAAACTTAAAAATCGAGTCGCAGCCCAAAATGCTAGAGACAAAAAGAAGGAAAGATCAGCAAAGATCGAGGATGTGATGCGCGATCTGGTGGAGGAGAACCGCCGGCTCCGCGCTGAAAACGAACGTCTTCGCCGTCAAAATAAAAATCTTATGAACCAGCAGAACGAGTCCGTCATGTATATGGAAGAGAACAACGAAAACTTGATGAACAGCAATGATGCATGCATCTACCAGAACGTCGTCTACGAAGAAGAAGTCGTCGGTGAGGTTGCACCAGTTGTCGTCGTCGGAGGAGAGGATCGCCGTGCCTTTGAATCAGCAGTGGGAACAGGCCCGATCCACCTCCATCAACAACAACATCAGCAACCAACTCCGTCGTATGGATTCCAAGAAGAACAACACAATCAGTGTGGATATGTATCTAACTATCATCTCGATTCTATGCAACCACATGGATCGCAACAAGAAGATGGACACCTCGAACAAATCCTCGAACATCTCAAGAGCCCAAGCGGAGAGTTCGATCGATTCGTTGCTGGCTACATTGAGGAAGGAGCAGACGGTTATGCAGCGTCTTGTTCAAGCGGATCCATGTACACATCTTCAGAAACGCGTGAAACACTTTCGCCGAATTCCCTAGCCATGTCCCCGTCGATGAGCAGCTCGAGCACTGACTGGGATGATGAGCTTTTGGGATGTGGAACCGAAACTGGAACTGGAACCGACGAGCTGCTTACCGACCCCGGAAACTGGAACTTTGAAACTTTCGACGAAAATTCAATCGACCTAAATTTCTTCCAAAATTAA

*unc-54 UTR* CATCTCGCGCCCGTGCCTCTGACTTCTAAGTCCAATTACTCTTCAACATCCCTACATGCTCTTTCTCCCTGTGCTCCCACCCCCTATTTTTGTTATTATCAAAAAACTTCTCTTAATTTCTTTGTTTTTTAGCTTCTTTTAAGTCACCTCTAACAATGAAATTGTGTAGATTCAAAAATAGAATTAATTCGTAATAAAAAGTCGAAAAAAATTGTGCTCCCTCCCCCCATTAATAATAATTCTATCCCAAAATCTACACAATGTTCTGTGTACACTTCTTATGTTTTTTACTTCTGATAAATTTTTTTGAAACATCATAGAAAAAACCGCACACAAAATACCTTATCATATGTTACGTTTCAGTTTATGACCGCAATTTTTATTTCTTCGCACGTCTGGGCCTCTCATGACGTCAAATCATGCTCATCGTGAAAAAGTTTTGGAGTATTTTTGGAATTTTTCAATCAAGTGAAAGTTTATGAAATTAATTTTCCTGCTTTTGCTTTTTGGGGTTTCCCCTATTGTTTGTCAAGATTTCGAGGACGGCGTTTTTCTTGCTAAAATCACAAGTATTGATGAGCACGATGCAAGAAAGATCGGAAGAAGGTTTGGGTTTGAGGCTCAGTGGAAG

*eat-4p* ATTTCTAATAAAACGGTCTACCATTTTGAGTCTATTATAGCCGAAAATCTCCAATGTGACTGTGACTTCTTAAAACTACTAAAACATTATTTGTCCATTTACATCTTCCTAAAACCGTATATCATCAAAAACATTCACAAAATCCGAAAAATGAGACAAAAATTTTTTTTTGATTGTTATTGCAATAAATCTAATAAAAATATTCATATATTGCCTGGCGCCCCCCATATCTCCATTTCCGGTCCCATCACCCCCACACCTCCAAGATTGATAGGTGGCTATAAGCATTTTTTGCATTTGAATGTGTTGCACCAGTAGTCATCATCATCATTATCTAAACTGACGTGATAGTAGGGGGCTTTCTAGAAGTCGATTTTCTATTAATGTCAACTTCATTCGTTGTCCCTTCCTTTCCCCGTCTTCCCTCACTTCCTTTTTTCTATTTTTTCCAGTGGTCCGTAGTGGGCGGCACCCGATTTTGACTTGAAATCAGACCCGTTTCCGGTTCTTTTGGTAGTTGTTAAGTTCTGATTCTATGACGTGGAGTGAAACAAAGAACGACCATATTTCATGTGTTGTGTTTTCTAGGCAGTCTAGGCAGGCAGTAGGCAGATACTGTCAAAATTGGAATATTTCCATCTTCTAAATACCCTCAACTATTTGTTAGCGCGTTTGAAACAATAATTGCAAAACATTTTTTTCGCATTGATTGGGGCATTTTGAAATTTAGAGTAAAATCGACTATCAACTGTCATTCCATAATAATTGGCAGAATTATTTTGGTTATGCCACCAAATAATCAATAAAGAAAGATTTCTGTCCTTATTAAACTAAAAATTGAAGCAACGGTAGAGTTGCTGAACACAGTCGCCGGCAAAATTTTTTAATTTTCTGCAAATTTGAAATTCTTTTGCGGGTTTTTAGTTATGGTTCTAATAGTGGTTAAAAGTCATTATAAAACACTTCAATTTTTTGTAATGCTTTCATTCGCAGTCGTGAAGTCGTGAAAACTCAGTTTTCACCTATCCTAACTTTGGAAGTCGTCCAAAAAATTATTTTAATTCTACAATTTTATATTTCTTTTTAAACATACGATGTGATCAATCCTACATCAATTCTGCAAATTT CTCACATTCTTGGAAGCTTGGTAATTTATCAGACTTTGACTGAAAATTTGAAAAAAAATTCAGTAATTTTTGGAGATCCCTTTAAATATTATTCTAGCATTGCCATATAGAATAATTGCAAAATTCAATTGGTTTCCTAGAAAGAAAATTAGATAATCTTATGAAAGAGAACCTAACCACAACAGGTGTTAAATATTGATTTAAACACTAAAAAAAGTCTCTCCTTCTACCTCTCTTCTCATAGCTTTCATGTCTTGAAGCTTTTCGGTTAATTCGAGCGGAAGACTTATCAAGGTACGTCATTTTCAATTACTTGTATACATCTCTCAGCTTCTTCTAATTTTCCATGTACATCTTAGATATTCTTTTTTCAGCGACTTGATATTAGGAAGTTTTGTGTTTTCAATTTTAAAGTTGGATTAATATAGAATACCAGTCTTTAAACACAAACCAACAAGGGTTCAATATCAAAATAGAAACCGAAAAAAAATATCGAAAACTGAAAAAATCAAATACATATCTAAAGCAAGCTATTCGAGAAATATTTATGCATTATAACAACTATGCAGCGCTTATATCTTTATTTTTCAACAAGTGTTCCAGCAACGAGAGTCTCTTCACCAAAAAGCCATCTATCAAAAACCAGGCAGTGAGTCCTAGAACCAAGCTTGTCAGAAGACAAGTGCATGTATAAATAAGTAGTAAAAGACGGGTGGGACCCAGCGCGGTGAGTAGTAGATCATAAAAAAGTGACTGAAAAGAAGGGGCGTTTCCTTTTCTTTATTACCTCTTCCTCTACTTCTTCTCATCTCAACTTGCTTTTTCCTTCCTCTTACAACAACTCCATCATCATCATCATCATTTTCAGAAACC

*tbh-1p* GAAATGTGTGTAGTGTCATGCTCAAAATAAAATCTGGCTATAGAGGTTACAAAAAGTATGGTTGCAATGATGTTACATAAATTGTGCCCACATGAAATTATGCAAGATAATTGGTAATGGGGTTGCTCATGTGATGACGGAAAATGCATAAATATATTTTGTTCACTTTTCCCCTATTCATGTGTGTAGAAGAATGTAAAAATACGCATGCATGTTTTGCATTTTTCAACATTTTTCGGCATTTTTGTACATTTTTCTACATCAGCCGACAGCCCACTTCACATGCTGAAGGCGAGAAACGCAGTGAGATGTGCCCCCGCAATGTCGAAAAATGCCGAAAAATGCAGAAACTGCATGCGTTTTTTTTACATTTTACTACACACATGGCTAGGGGAAAAAAAAATGTAAAAATACATTTTTGCAATTTTCGTAATAACATAAATAACCCCATAAATGTCTTTTTTATAACTAGAATTCTAGAATTTCCTTAAAGCCACAAAGATATTCCTCTGAAATTTTATTTTTATCTGACTGTGACTTATCTTGTTTCTATTATGTGCATTTTTTGTCGTTAGCTTTGTCGGCTCCAAATTAAATTTCATAAATTATCAATAAATTATCATTGCACAAATATGTAACAGTATGCAATATTCACATTACAATTTTATTTTGATTTCTCTACTGGTAGAGTTGATGGTTGCGTTGGTGCAGCTGCAGTAGTTCTTACATATTTTGAAGAGCGAAGAGCCTCGGCGGATAAAAAGTAGTGTCCATTTTCTCGTCGGATACTCCGGAGCAGCGCAGTTTGATGTATCAGCTTCTCTGCTGGACATGTTTGAATCTAAATTTTCAGAACTTTCAAAAGAACTAACAATTTTCATATCCCACCAGCTTTTGACGTTTTCCAATCTCGAGCTCTTGAAACAAATTCCTATATGTTGTTTGCGTGTCAAGCGGCACATTTGATCTGAAAAATGAAATTCTTGATAAGAATTCAATTATCATGGAATGCTTCATTACACTGGCAAAGCGGATGCTTTTCTCGAAAGGCGTGTTCTGGCGACGGATGCATCAAATGGTTGATATTGGCTACGGTGGGTGGTTTCAGAATAAAAGGGGATTTTCGGAGCATCTCCTGTTTCAGATTCAGTTATTTCTCTGAAAATAAAAACTTATCAACTTTTTCATTTCATGTGTAAGATTTGGTGGTTATCAGGATCAATACTATAAAATCTTAAAAAAAGTAAATTTATTTTTATTTTTTAAATATTTCCGAAAAATAGTTTTTTGTTGAGTGGTTTTTCACAACCGCGATGTCTCTTCATTTTATTACACAAGGTATCACTTCTTTCCTTTCTTGTGTTTTTATGACTAGCATATTCTTTTTTTGAAAGATGTGCAGTTGACAGTTTTCAAATACTCCCTGTCATTGTCACTTGTCATTTTCATACCTCTTCTCCCTTGCGTGCGGCACTTTCCACGTATAAGCGGCGTTGTTTGTGGTGCGCCCGTAGA

*unc-25p* TCCTACCTTTTCTATCCCACTTTTTCCCCGGAATTTTGAGATTTGACATGATTTTTCAGAGAATTTCGAGTTTTGAGAAAAAAAAAATCAAAAAAGCGATTTTTTGGTCAAAAGCCGAAATTTAAAGCTAGTTTTTTTGGAAATATCGATTTTTTTTTTGAGAATTTTTCGATTTTCCAAAAAAAAAATCGAAAGTTGTTTCCCGAAAAGCCGAAATTTTTTTTGAGATATCGAAAAATCGGAGAATTTTTCGATGTTTCCCAGAAATACTAAAATTTTGAATTTTCCGGAAAAAAAATCGTTAAAAAATTAGTTTCTTTTAAAGACCGAAATTTAAAGTTTTTTTTTTGGAAATATCGATTTTTTCCGAGATATCTAAAATTGAAAGAAATTTTCGATTTTCTGAGAAAAATACGAAAAAATCGAAAAAAAAACCCAAATTTCGGTTTTTTCGTAAAAAAAGTCGTAAAAATGTAATTTTTTTCCTGAAAAATCGGAATTTTTTA CAAAATATCGGAAAATACTCAAAAAAAAGCTGAAAATTTCGATTTTCCGGAGAAAAAATCTTTTTAAAAAAATATTTTTTTTTTCAGAAAATAAGAAAAGCCGAAATTTATAACTATTTCTCCGGAAATTCGAAATTTTTAACGAAATATCGGAATAATTTTTAGATTTTTCAAGTTTGACTTTGCGAGAAAAAAATCGGAAAAACCTCGATTCGACGCCGAAAAATGCTCCTTTTCGAAAAGATTTTTGAAATTTCAGAAAATCGACATGCAAGCGCGCTCCACGGCGAAATGACAACGATGATCCACCGCCCTCAAAAAGTTGGGTCTCGTTAGGTATTTGGCGGTAAAACTGGTAAAACTCCAGTTTTGCCTCCAACGAGACCCAATTTTTTGGGGCGGTGGTGGAGCGCGCTTGCACAAGCTGAAAGCATTTTTCTGCGACTCGATAATATTTTGAAAACCTGTGTCAATTCTCGAAATTCTTTTTTTAAAAAATAATCCCGAGCTTCTCTCAGTCCTCCTCTATGAGGATGTTCCTTTTTTTTGGTTTTTCAATTTTTTTTAAAATTCCAAATTTCTGTTGTGCAATTCACTTCCCCCCAAGAAATCCCAAAAATCCCCAGTTTTCCCCAAAAATGTTCCGTTTTCATGTGATTTTTCCCCCATTTTTAAAACATTTTTTTGACTTTTTTTTAAAATGATTATTATTATTGTTTTCTATTTCATGGCCGGTAAATTATTTTTTTTCTTTCTTTTTTTTTGCTCTTTTTTTTCAAGAATTTTCGAATTGTTTGAAGGGCTGCTCATCTAATCTTTTGTCATTTTGTTCTGATGCCATCATTTCTGAGAGGACCTTTGAAGACTCGTCACGAAACGGGAGGGGGGCTCAAGTGAGCATTATTATTATTATTATTGTCGCAAAAAGTTTACCCCGGGCTCCCCCTGGCTCCCCTCTTTGAGCAAGGGTTTAAGGGCTCATTTTGATGACGAATTGCTCATTGGGATTATAGTCACGCCCCTCTTTTGGAGCAACTACACAACTGAGCCACAGTAATCCTTGGGGGCGGGGTCAGTAGGACCCCCTCCGGAATAGGGAAAAGCTCAGTTCACCGCCAAA A

### Neuron Count

Worm Atlas (https://www.wormatlas.org/neurotransmitterstable.htm) summarizes the proposed neuron location for glutamatergic, octopaminergic, and GABAergic neurons in hermaphrodites.

| Neuron Type | Proposed Number | Location by Body Region |
| --- | --- | --- |
| Glutamatergic | 79 | Head: ADA, ADL, AFD, AIB, AIM, AIZ, ASE, ASG, ASH, ASK, AQR, AUA, AWC, BAG, FLP, IL1, OLL, OLQ, RIA, RIG, RIM, URY<br>Pharynx: M3, M1, I2, I5<br>Ventral nerve cord and body: ALM, AVM, CP0, CP5, CP6, CP7<br>Tail: DVC, LUA, PHA, PHB, PHC, PLM, PVD, PVQ, PQR, PVR |
| Octopaminergic | 2 | Head: RIC |
| GABAergic | 32 | Head: AVI, DD1-6, DVB, RIB, RIS, RME, SMDD/V, VD1-13<br>Ventral nerve cord and body: DD2-4, VD3-11<br>Tail: DD6, DVB, VD12-13 |

### C. elegans microscopy

#### Stereoscope

For whole-worm imaging of *vha-6p::Q40::YFP*, DHS-3::GFP, and VIT-2::GFP strains, synchronized animals were grown on RNAi plates seeded with EV bacteria or RNAi bacteria. Animals were imaged at day 1 of adulthood and for aging experiments also imaged at day 5 and day 9 of adulthood. 10+ animals were placed in a pool of 100 mM sodium azide in M9 on standard NGM plates without bacteria to induce paralysis. Paralyzed animals were then lined alongside each other and imaged on a Leica M205FCA automated fluorescent stereomicroscope running LAS X software and equipped with a standard GFP filter, Leica LED3 light source, and Leica K5 camera. For all imaging experiments, 3 biological replicates were performed with 2 technical replicates each, and 1 representative image was chosen for use in figures. For *vha-6p::Q40::YFP* quantification, Fiji^14^ was used to draw a region of interest along the posterior half of each group of worms, and integrated density was measured. Graphing and statistical analysis were performed with GraphPad Prism 10 software using a Mann-Whitney test unless otherwise stated. *vha-6p::Q44::YFP* quantification was performed by counting the number of protein aggregates in individual worms and statistical analysis was performed with GraphPad Prism 10 software using a Mann-Whitney test.

#### Widefield and confocal imaging

Widefield imaging utilized a Leica THUNDER Imager equipped with a 63x/1.4 Plan AproChromat objective, standard dsRed filter (11525309), Leica DFC9000 GT camera, a Leica LED5 light source, and run on LAS X software. For high-resolution imaging of DHS-3::GFP and ER morphology, imaging was performed using a Leica Stellaris 5 confocal microscope equipped with a white light laser source and spectral filters, HyD detectors, 63x/1.4 Plan ApoChromat objective, and run on LAS X software. Animals were placed in 100 mM sodium azide solution on a glass slide to induce paralysis and imaged within 5 minutes of slide preparation to prevent artifacts from prolonged exposure to sodium azide. For imaging experiments, 3 biological replicates were performed with 1 or 2 technical replicates each, and 1 representative image was chosen for use in figures.

#### Intestinal bacteria invasion assay

Assessing intestinal bacteria invasion was performed as previously described^15^. Animals were L1 synchronized via bleaching and plated on RNAi plates containing a bacterial lawn derived from a mixture of 80% HT115 bacteria containing EV and 20% HT115 bacteria expressing mCherry. Once at the desired age, animals were manually transferred onto a standard OP50 plate and allowed to feed on OP50 for 2 hours at 20°C to facilitate clearance of mCherry bacteria. For imaging, worms were paralyzed by exposure to M9 solution containing 100 mM sodium azide and arranged on a standard NGM plate without bacteria. Images were captured using a Leica M205FCA fluorescent stereomicroscope equipped with a standard dsRed filter as described above. For each of the 3 biological replicates, 2 technical replicates with 13 animals per replicate were performed and 1 representative image was used for figures. The percentage of animals exhibiting bacterial invasion was quantified and plotted with GraphPad Prism 10 software for each technical and biological replicate. Statistical analysis was conducted across all replicates using a Mann-Whitney test with GraphPad Prism 10 software.

#### ER Secretion Assay

Assaying of ER secretory function was performed as described previously^16^. Transgenic control animals and *eat-4p::xbp-1s*, *tbh-1p::xbp-1s*, and *unc-25p::xbp-1s* animals expressing VIT- 2::GFP were bleached to obtain eggs. Eggs were then placed on glass slides and imaged using a Leica THUNDER Imager equipped with a 63x/1.4 Plan AproChromat objective, standard GFP filter (11525309), Leica DFC9000 GT camera, a Leica LED5 light source, and run on LAS X software. Images were quantified using Fiji and drawing a region of interest around each individual egg to obtain an integrated density value. 4 independent biological replicates were performed. SuperPlots^17^ were created using GraphPad Prism 10 software where large dots represent the median value of each biological replicate and small dots represent single eggs with different intensities of colors representing eggs from the same biological replicate; lines indicate median and interquartile range. All statistical analyses were conducted using Mann– Whitney testing with GraphPad Prism 10 software. For whole worm imaging, animals were raised to day 3 of adulthood and imaged using the Leica M205FCA fluorescent stereomicroscope equipped with a standard GFP filter as described above.

#### Oil Red O Staining

Oil Red O fat staining was performed as previously described^18^. Briefly, worms were bleached, and eggs were plated to obtain a synchronous population. Worms were grown on RNAi plates with a lawn of EV bacteria and aged to day 3 of adulthood. Aging was performed in the absence of FUDR by gravity settling in M9 solution and aspirating to remove progeny. For staining, worms were washed off plates using a PBS + 0.01% Triton solution, rocked for 3 minutes in 40% isopropyl alcohol, pelleted, and then stained with Oil Red O in diH_2_O for 2 hours while rocking at room temperature. Worms were pelleted and washed in PBS + 0.01% Triton for 30 min before being imaged at 20× magnification with a Leica THUNDER Imager Flexacam C3 color camera and run on LAS X software. To quantify somatic fat depletion, worms were scored as previously described^19^. The level and distribution of fat was placed into categories of non-somatic lipid depletion, displaying no loss of fat and being darkly stained throughout the body, and somatic lipid depletion, being stained largely in the germ cells. At least 100 worms were scored for each condition over 3 biological replicates.

### *C. elegans* RT-qPCR and RNA-seq analysis

For collection of RNA, we used *glp-4(bn2)* animals to eliminate progeny. After bleaching and L1 arresting, all animals were raised at 22°C (the restrictive temperature for our backcrossed *glp- 4(bn2)* strain) for 3 days to collect animals at day 1 of adulthood. Approximately 1000 animals were used per condition. Worms were collected using M9 and transferred to TRIzol solution and underwent 3 freeze/thaw cycles between liquid nitrogen and a 37°C bead bath with a 30-second vortexing step between each cycle to lyse worms. Following the final thaw, chloroform was added at a ratio of 1:5 chloroform/TRIzol, and aqueous separation of RNA was achieved by centrifugation using a heavy gel phase-lock tube (VWR, 10847–802). The aqueous phase was mixed with isopropanol at a 1:1 ratio and applied to a QuantaBio Extracta Plus RNA kit (95214) for RNA purification according to the manufacturer’s instructions.

Library preparation and sequencing was conducted at Novogene using their standard pipeline using paired-end, polyA selection, first-strand synthesis, and an Illumina NovaSeq6000. Each condition was measured with 3 biological replicates. Gene expression levels were quantified using STAR-2.7.3a^20^ with WBcel235 as the reference genome. Fold changes were determined using DESeq2^21^. Gene targets of XBP-1 were defined based on previous experimental findings^22^. Gene Ontology (GO) enrichment analysis was performed using WormEnrichr^23,24^. *rgef-1p*, *dat-1p*, *tph-1p*, *eat-4p*, *tbh-1p*, and *unc-25p* driven expression of *xbp-1s* was compared to the N2 wild-type control.

For RT-qPCR, cDNA synthesis was conducted using qScript cDNA SuperMix (QuantaBio, 101414-102) with 500 ng of RNA. RT-qPCR was performed using NEB Q5 DNA polymerase following the manufacturer’s guidelines and utilizing the primers listed below. Each condition was assessed using 3 biological replicates. QuantStudio 3 (Thermo Fisher) was used for quantification using a standard curve method.

### Lifespan measurements

For lifespan experiments, animals were grown on RNAi plates on either EV or RNAi bacteria from L1 stage. At day 1 of adulthood, animals were washed off plates with M9 solution and then moved to plates containing 100 µL of 100 mg/mL FUDR to eliminate progeny. One replicate of lifespan assays was performed using a lifespan machine^25^ with others being done by hand. Tunicamycin survival assays were performed based on established protocols^26^. For tunicamycin assays, animals were moved onto plates supplemented with 25 µg/mL tunicamycin in DMSO directly in the plate. Animals were grown at 20°C and checked every 2 days for viability. Animals were considered dead if they did not exhibit any movement when prodded with a platinum wire at both the head and the tail. Animals that exhibited bagging, intestinal leakage, desiccation on the side of the plate, or other deaths unrelated to aging were scored as censored. All lifespans were performed on >3 biological replicates. Lifespan assay survival curves were plotted using GraphPad Prism 10 software and statistics were performed using a Log-Rank test in GraphPad Prism 10. Representative data are depicted in figures and a table of all lifespan assays performed is available in **Table S7**.

### Caenorhabditis elegans brood size assay

Brood assays were measured as previously described^26^. Bleaching was used to obtain a synchronized population of animals, and 10 L4 stage animals were transferred onto individual plates. Every 12 hours, the animals were moved onto new plates, while plates containing eggs were stored in a 15°C incubator for 2–3 days. All surviving progeny on each egg-laying plate were counted and totaled to determine the brood size. SuperPlots^17^ were created using GraphPad Prism 10 software where large dots represent the median value of each biological replicate and small dots represent single animals with different intensities of colors representing animals from the same biological replicate; lines indicate median and interquartile range. All statistical analyses were conducted using Mann–Whitney testing with GraphPad Prism 10 software.

### Caenorhabditis elegans thrashing assay

Thrashing assays were conducted on animals synchronized via bleaching and aged on RNAi plates containing FUDR from day 1 of adulthood. Upon reaching the desired age, plates containing adult animals were flooded with 100 μL of M9 solution, and 30-second videos were recorded using an M205FCA stereomicroscope equipped with a Leica K5 microscope running LAS X software. Thrashing movements were manually recorded over a 10-second period. A bending of more than 50% of the animal’s body in the opposite direction was deemed a single thrash. Representative data from 3 independent biological replicates are presented.

SuperPlots^17^ were created using GraphPad Prism 10 software where large dots represent the median value of each biological replicate and small dots represent single animals with different intensities of colors representing animals from the same biological replicate; lines indicate median and interquartile range. All statistical analyses were conducted using Mann–Whitney testing with GraphPad Prism 10 software.

### Fast kill assay

Fast kill assays were performed as previously described with minor modifications^27,28^. *Pseudomonas aeruginosa* (PA14) cultures were grown overnight at 37°C for 14-15 hours. 5 μL of overnight culture was spread over 3.5 cm peptone glucose media plates (1% Bacto-Peptone, 1% NaCl, 1% glucose, 1.7% Bacto Agar) containing 0.15 M sorbitol, using a spreader made from an open loop tipped glass pasture pipette. The plates were incubated at 37°C for 24 hours and then at 25°C for 48 hours. Following this, 30 to 40 synchronized L4 animals were placed on each plate. Assays were performed at 25°C. Survival of animals was plotted over a period of 8 hours with intervals of 2 hours. An animal was deemed dead when it no longer responded to touch. 3 biological replicates with 3 technical replicates were performed for a total of 9 replicates. Survival rates were measured wherein 100% survival was indicated as an integer “1” and the fraction of survival populations at every time point was represented as decimal values.

### Forced food choice

Forced food choice assays were performed as described previously^29,30^. Plates were made using the same recipe for NGM plates aside from the addition of 0.35% peptone. A single colony of *Pseudomonas aeruginosa* (PA14) and *E. coli* OP50 bacteria were inoculated into separate 3 mL of LB for overnight primary culture at 37°C. The following day, the OD_600_ of each of the cultures was diluted to an OD_600_ of 1.0. A culture of PA14 was transferred as a line along the center of an NGM plate using a glass Pasteur pipette bent at a 90-degree angle. Next, 15 μL of OP50 culture was seeded as a dot onto the plate 2.5 cm away from the center and 0.5 cm away from the edge of the plate. The plates were dried and transferred to 37°C for 24 hours, followed by incubation at 25°C for 48 hours. On the day of the assay, the plates were removed from the 25°C incubator and allowed to reach room temperature. Worms were washed three times in M9 solution before being placed onto the assay plate diametrically opposite to the OP50 dot at 2.5 cm away from the center. The proportion of worms found on or off each food was recorded after 1 hour, 2 hours, 4 hours, 6 hours, and 8 hours. After each time point, the population was scored in which -1 represented 100% of the population on PA14 and +1 represented 100% of the population on OP50. The movement index was then calculated towards the OP50 dot using the formula:

<u>(“A” population of worm on OP50 dot – “B” population of worms on PA14 line)</u> (“A” population of worm on OP50 dot + “B” population of worms on PA14 line)Each assay was done in biological triplicate with technical triplicates for a total of 9 replicates. Statistical analyses were conducted using Mann–Whitney testing with GraphPad Prism 10 software.

### Statistics

All statistical analyses were performed using GraphPad Prism 10 software. No assumptions were made about data distribution. Mann-Whitney testing was used for most comparative analyses with p-values less than 0.05 considered significant unless otherwise specified. To determine statistical significance of lifespan data, a log-rank test was performed with p-values less than 0.05 considered significant. At least 3 biological replicates were performed for each experiment unless otherwise noted.

## Results and discussion

### Overexpression of xbp-1s in glutamatergic, octopaminergic, and GABAergic neurons

In previous studies, serotonergic, dopaminergic, and RIM/RIC neurons have been identified to be involved in neuron-to-body communication of UPR^ER 3,4^. However, these four neuron subtypes make up only ∼18 of the 302 neurons in *C. elegans*, raising the question of what other neuronal subtypes may be involved in neuron-to-body UPR^ER^ communication. Indeed, transcriptomic analysis of previously published datasets^3,8^ reveals only a minor overlap of differentially expressed genes between worms expressing *xbp-1s* in dopaminergic and serotonergic neurons as compared to pan-neuronal expression (**Fig. 1A-C**). In addition, the expression levels of differentially expressed genes under pan-neuronal *xbp-1s* expression are largely dissimilar when *xbp-1s* is expressed in serotoninergic or dopaminergic neurons, or both concurrently (**Fig. 1D**). This strongly suggests that other neuronal subtypes are involved in non- autonomous UPR^ER^ signaling.

**Fig. 1.**
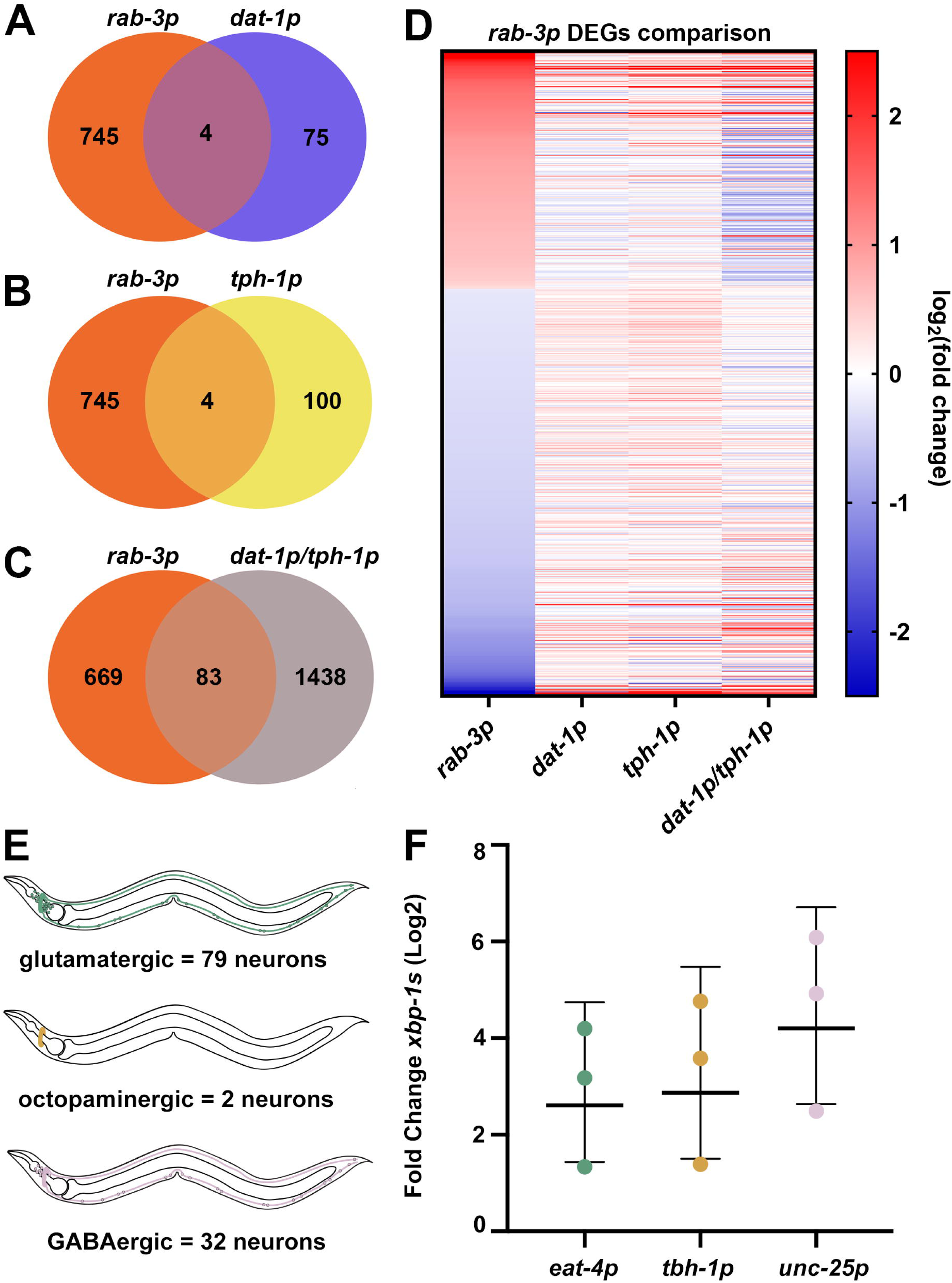
Dopaminergic and serotonergic *xbp-1s* together do not recapitulate pan-neuronal *xbp-1s* overexpression. A comparison of differentially expressed genes (p-value ≤ 0.01) between worms expressing *xbp-1s* pan-neuronally (*rab-3p*), and in either **(A)** dopaminergic (*dat- 1p*) neurons, **(B)** serotonergic (*tph-1p*) neurons, or **(C)** concurrently in both dopaminergic and serotonergic (*dat-1/tph-1p*) neurons. For a complete list of differentially expressed genes see **Table S3**. For a complete list of genes represented in Venn Diagrams, see **Table S4**. **(D)** Heat map of differentially expressed genes in worms expressing *xbp-1s* pan-neuronally with corresponding expression levels in serotonin, dopamine, and both serotonin and dopaminergic *xbp-1s* expressing animals. Warmer colors indicate increased expression, and cooler colors indicate decreased expression. See **Table S5** for a list of gene names and expression values. Schematic of each neuronal type explored in this study: glutamatergic, *eat-4p* (n = 79, location: head, pharynx, ventral nerve cord and body, tail), octopaminergic, *tbh-1p* (n = 2, location: head), GABAergic, *unc-25p* (n = 32, location: head, ventral nerve cord and body, tail). qPCR of transcripts of glutamatergic *xbp-1s* (green), octopaminergic *xbp-1s* (yellow), and GABAergic *xbp-1s* (pink) animals grown on empty vector from hatch. RNA was isolated in day 1 adults and data are compared using a standard curve with data represented as relative fold change against control. Each dot represents a biological replicate averaged across three technical replicates per sample. Lines represent geometric mean with geometric standard deviations.

Previously, we performed a screen of several neurotransmitter signaling pathways involved in neuronal communication of UPR^ER^, which revealed glutamatergic, octopaminergic, and GABAergic neurons as candidates involved in this signaling event^3^. Glutamate is a widely utilized, excitatory neurotransmitter in both invertebrate and vertebrate systems^33^; octopamine is a *C. elegans*-specific neurotransmitter similar to the mammalian norepinephrine, and is involved in immune response^34^; and gamma-aminobutyric acid (GABA) is a widely utilized neurotransmitter that has been found to function as both an excitatory and an inhibitory signal in *C. elegans*^35^. *C. elegans* possess 79 glutamatergic neurons, 2 octopaminergic neurons, and 32 GABAergic neurons in hermaphrodites (**Fig. 1E**) (Loer CM, Worm Atlas, https://www.wormatlas.org/neurotransmittercriteria.htm).

To determine the potential involvement of glutamatergic, octopaminergic, and GABAergic neurons in neuron-to-body communication of UPR^ER^, we overexpressed *xbp-1s* in each neuronal subtype using the *eat-4* promoter for *xbp-1s* overexpression in glutamatergic neurons^36^ (hereafter referred to as glutamatergic *xbp-1s*); *tbh-1* promoter for *xbp-1s* overexpression in octopaminergic neurons^37^ (hereafter referred to as octopaminergic *xbp-1s*); and the *unc-25* promoter for *xbp-1s* overexpression in GABAergic neurons^38^ (hereafter referred to as GABAergic *xbp-1s*). We confirmed by quantitative PCR (qPCR) that all three subtypes display an increase in *xbp-1s* mRNA, although our data did not reach statistical significance (**Fig. 1F**).

### Glutamatergic, octopaminergic, and GABAergic xbp-1s alter distinct transcriptional pathways

To more thoroughly investigate the impact of neuronal subtype UPR^ER^ on the periphery, we performed whole-worm RNA sequencing on animals overexpressing *xbp-1s* in glutamatergic, octopaminergic, and GABAergic neurons. Glutamatergic and octopaminergic *xbp-1s* resulted in sizable changes to gene expression, while more mild changes occurred with GABAergic *xbp-1s* (**Fig. 2A-C**). Interestingly, the majority of differentially expressed genes were unique to each condition, suggesting distinct responses were induced by each neuronal subtype (**Fig. 2D**). This adds more insight into a previous study that identified distinct pathways activated downstream of serotonergic and dopaminergic *xbp-1s* ^3^.

**Fig. 2.**
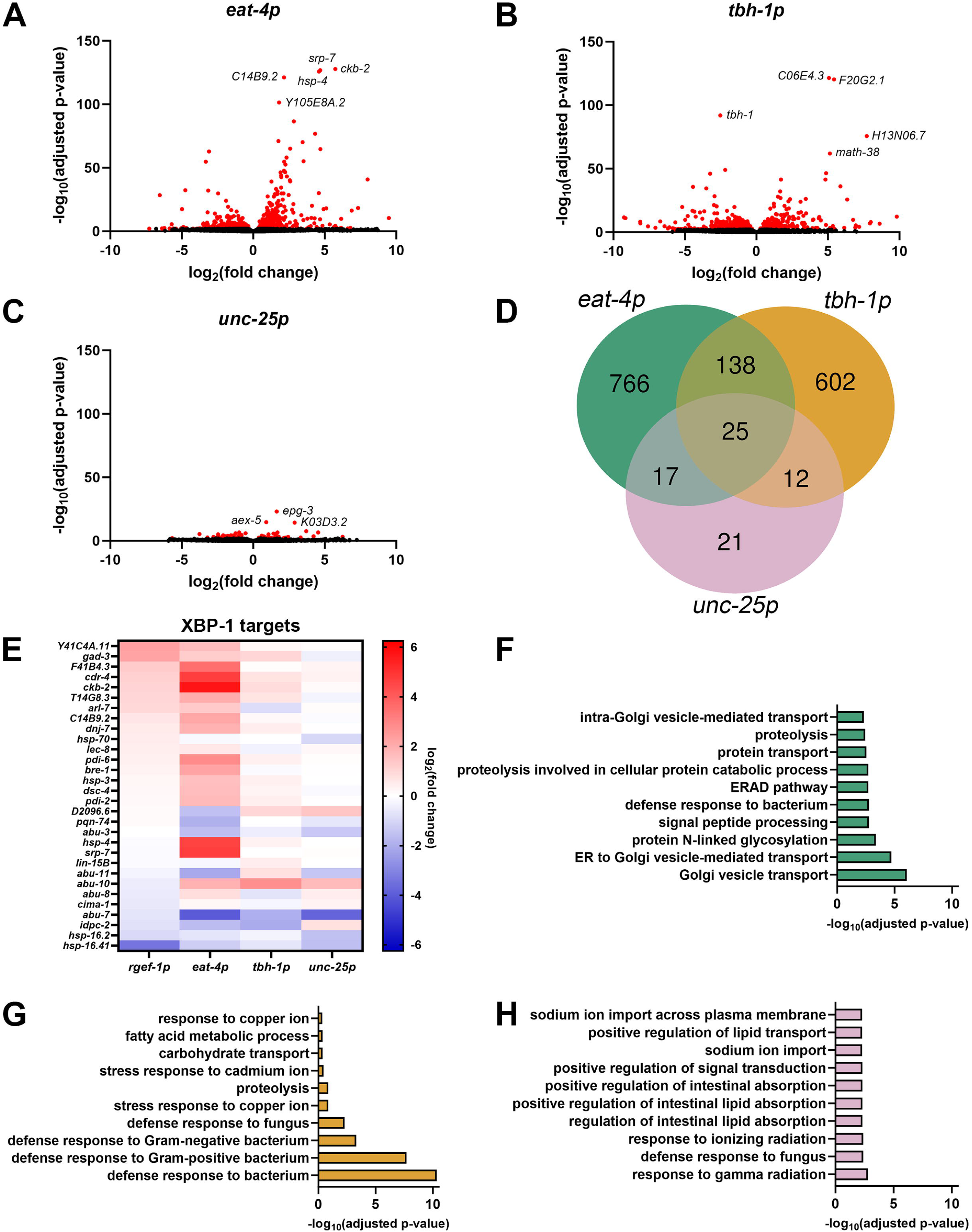
Glutamatergic, octopaminergic, and GABAergic *xbp-1s* modulate distinct transcriptional pathways. Volcano plots of whole-body genome-wide changes in gene expression upon *xbp-1s* overexpression in **(A)** glutamatergic, **(B)** octopaminergic, and **(C)** GABAergic neurons. Red dots indicate significantly differentially expressed genes with p-value ≤0.01. See **Table S3**. **(D)** Comparison of differentially expressed genes (p-value ≤ 0.01) between worms expressing *xbp-1s* in glutamatergic, octopaminergic, and GABAergic neurons. For a complete list of differentially expressed genes see **Table S4**. **(E)** Heat map of XBP-1s target gene ^31^ expression under neuronal, glutamatergic, octopaminergic, and GABAergic *xbp-1s*. Warmer colors indicate increased expression, and cooler colors indicate decreased expression. See **Table S5**. Top ten most enriched gene ontology terms of differentially expressed genes upon *xbp-1s* overexpression in **(F)** glutamatergic, **(G)** octopaminergic, and **(H)** GABAergic neurons. See **Table S6**.

To further characterize the similarities and differences between peripheral response to neuronal subtype UPR^ER^, we directly compared our glutamatergic, octopaminergic, and GABAergic *xbp- 1s* animals to previously published RNA-seq datasets ^3,8^. First, we sought to determine the overlap between neuronal subtype *xbp-1s* overexpression with pan-neuronal *xbp-1s* overexpression (hereafter referred to as neuronal *xbp-1s*), as we would expect that neuronal *xbp-1s* includes each neuronal subtype. We compared neuronal *xbp-1s* using two different promoters, *rab-3p* and *rgef-1p* and were surprised to find that while there was significant overlap between these two neuronal *xbp-1s* strains, a majority of differentially expressed genes were not shared (**Fig. S1**). Since this could potentially be due to leakiness of the *rab-3p* compared to the *rgef-1p*^39,40^, in our subsequent studies, we focused on making comparisons to results from the *rgef-1p::xbp-1s* strain (which we will continue to refer to as neuronal *xbp-1s*).

As expected, neuronal *xbp-1s* animals display altered expression of a large number of direct XBP-1s targets^31^. Interestingly, we see that glutamatergic *xbp-1s* induces many of these same XBP-1s targets and to an even greater extent than neuronal *xbp-1s* (**Fig. 2E**). These data suggest that glutamatergic *xbp-1s* activates a more canonical UPR^ER^ signature involved in conventional protein processing pathways. Gene ontology (GO) enrichment analysis supported this idea, as the most enriched biological processes included pathways related to ER function and protein homoeostasis, including ER to Golgi vesicle-mediated transport, protein N-linked glycosylation, endoplasmic-reticulum-associated protein degradation (ERAD) pathway, and proteolysis (**Fig. 2F**). However, a majority of differentially expressed genes in glutamatergic *xbp- 1s* are still distinct from neuronal *xbp-1s* (**Fig. S2A**), suggesting that these protein homeostatic pathways are being regulated in different ways in each condition. Interestingly, glutamatergic *xbp-1s* transcriptionally regulates an entirely different set of genes than serotonergic *xbp-1s* (**Fig. S2E**), although these animals were also shown to induce canonical protein homeostasis pathways^3^. Altogether, these data show that even amongst neuronal subtypes that share a similar peripheral response (e.g., protein homeostasis), the specific genes targeted in these similar pathways are distinct, highlighting the fact that non-autonomous UPR^ER^ is dramatically different based on which neuronal subtype is involved.

Octopaminergic *xbp-1s* showed smaller gene expression changes to XBP-1s targets in comparison to glutamatergic *xbp-1s*, being more reminiscent of the levels found in neuronal *xbp-1s* (**Fig. 2E**). However, similar to glutamatergic *xbp-1s*, when all differentially expressed genes for octopaminergic *xbp-1s* were compared to neuronal *xbp-1s*, the majority of differentially expressed genes were distinct (**Fig. S2B**). The differentially expressed genes identified were entirely different from those found in serotonergic and dopaminergic *xbp-1s* (**Fig. S2D-E**). GO analysis identified that the most dramatic changes in gene expression in octopaminergic *xbp-1s* were defense response pathways, particularly those involved in immune response (**Fig. 2G**). These data are consistent with previous findings that showed pathogen response in *C. elegans* is associated with UPR^ER^ induction^41^ and a role for non-autonomous signaling in this response^11^, potentially through octopaminergic signaling^34^. These data add an additional downstream function of non-autonomous UPR^ER^ in regulation of immune response, potentially downstream of octopaminergic neurons.

Finally, GABAergic *xbp-1s* activation caused minimal changes in gene expression overall, with very little overlap with other neuronal subtype *xbp-1s* (**Fig 2E, S2C-E**). Although the gene expression changes were minor, GO analysis did reveal some pathways previously associated with UPR^ER^ induction, including lipid remodeling^3,5^ (**Fig. 2H**). When looking at all genes related to canonical UPR^ER^ (**Fig. S2F**) and XBP-1s targets (**Fig. S2G**), glutamatergic *xbp-1s* animals displayed the most significantly differentially expressed genes, while octopaminergic *xbp-1s* had more subtle effects and GABAergic *xbp-1s* displayed no major differences. Interestingly, octopaminergic *xbp-1s* also had significantly differentially expressed genes for the UPR^MT^ (**Fig. S2H**), which further adds evidence to its role in immune response as UPR^MT^ activation has been directly linked to response to pathogens^42^. Glutamatergic *xbp-1s* animals also displayed significantly differentially expressed genes for the heat-shock response (HSR, **Fig. S2I**) and oxidative stress response (OxSR, **Fig. S2J**), which is consistent with previous reports that suggested some overlap between UPR^ER^ and the HSR ^43^ and OxSR ^44^. Finally, octopaminergic *xbp-1s* animals show significantly differentially expressed genes for genes related to translation (**Fig. S2K**), another feature often correlated with UPR^MT^ activation^45^. Altogether, our data adds more evidence to the previously proposed model^46^ that specific neuronal subtypes participate in activation of unique downstream pathways in response to stress.

### Glutamatergic, octopaminergic, and GABAergic xbp-1s do not alter general organismal health, and only octopaminergic xbp-1s is sufficient to extend longevity

Next, we sought to test the impact of neuronal subtype *xbp-1s* on general organismal health, as previous studies have shown that neuronal *xbp-1s* results in a significant improvement in longevity and animal health, with a reduction in reproductive health ^2,37^. Interestingly, we found that octopaminergic *xbp-1s* animals had a bimodal population of lifespan (**Fig. 3A**), which correlates with animal size (**Fig. 3B**). Specifically, a proportion of octopaminergic *xbp-1s* animals display a stunted growth phenotype, and these animals tend to have a mildly reduced lifespan compared to control animals. In contrast, animals that display regular size display a significant lifespan extension. Since the reduction in lifespan for the stunted growth animals can be due to a number of pleiotropic and unrelated reasons, we opted to perform all further analyses on octopaminergic *xbp-1s* animals on the long-lived, “normal” sized animals.

**Fig. 3.**
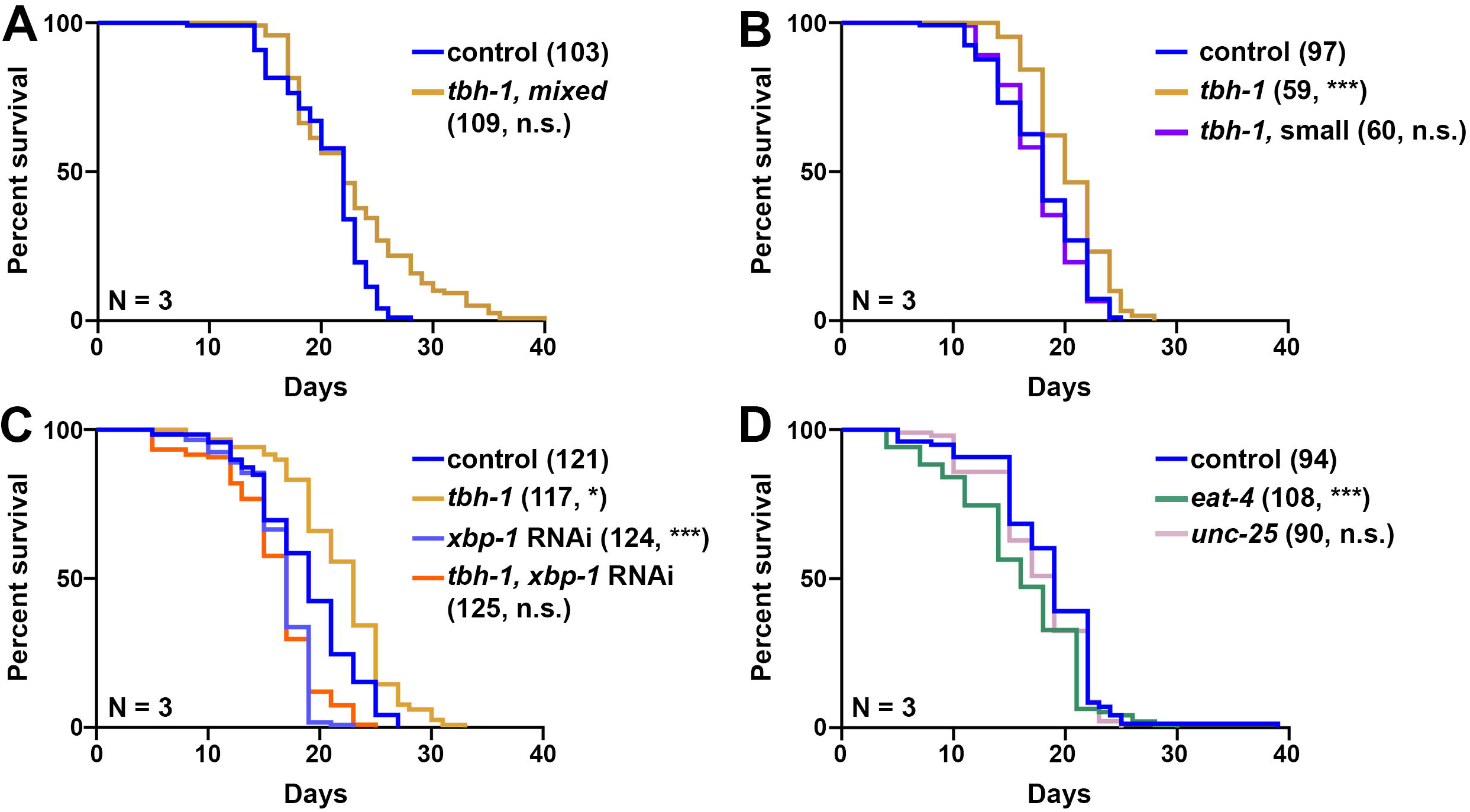
Octopaminergic *xbp-1s*, but not glutamatergic or GABAergic *xbp-1s*, is sufficient to extend lifespan. **(A)** Lifespan measurements of control (blue) and a mixed population of both “normal” and “stunted” growth octopaminergic *xbp-1s* (yellow, *tbh-1p*, mixed) animals. **(B)** Lifespan measurements of control (blue) and octopaminergic *xbp-1s* animals. Octopaminergic *xbp-1s* animals were separated into normal size (yellow, *tbh-1p*) and stunted growth (purple, *tbh-1p,* small). **(C)** Lifespan measurements of control (blue, light blue) and octopaminergic *xbp- 1s* animals (*tbh-1*, yellow, orange) grown on either EV or *xbp-1* RNAi. **(D)** Lifespan measurements of control (blue, light blue), glutamatergic *xbp-1s* animals (*eat-4p*, green), and GABAergic *xbp-1s* (*unc-25p,* pink) animals. Lifespans were scored every 2 days and data is representative of 3 biological replicates (N). Sample size (n) is written next to each condition followed by significance measured using Log-Rank testing: n.s. = not significant, * = p < 0.05, *** = p < 0.001. All statistical analysis is available in **Table S7**.

Importantly, this lifespan extension of octopaminergic *xbp-1s* animals was fully dependent on *xbp-1* (**Fig. 3C**), similar to all other neuronal *xbp-1s* paradigms previously established^2,3^.

Interestingly, glutamatergic or GABAergic *xbp-1s* animals did not display any increase in lifespan, and glutamatergic *xbp-1s* animals actually had a mild decrease in lifespan (**Fig. 3D**). In addition, while we saw a mild decrease in brood size in glutamatergic, octopaminergic, or GABAergic *xbp-1s* animals, these differences were not statistically significant (**Fig. S3A-C**).

Finally, general organismal health was also unchanged as no change in motility was observed, except a mild but statistically not significant increase in day 1 thrashing of octopaminergic *xbp- 1s* animals (**Fig. S3D-F**). Thus, only octopaminergic *xbp-1s* was sufficient to promote longevity, with only minor – if any – changes in other healthspan metrics.

### Glutamatergic, octopaminergic, and GABAergic xbp-1s improve immune function

To further investigate a potential mechanism whereby octopaminergic *xbp-1s* animals promote longevity, we next measured common features of UPR^ER^ induction. Our transcriptomics analysis revealed that glutamatergic, octopaminergic, and GABAergic *xbp-1s* displayed significant changes in immune response related genes (**Fig. 4A**), with octopaminergic *xbp-1s* animals having defense response against bacteria as one of the most significantly enriched GO terms (**Fig 2G**). Therefore, we measured the impact of *xbp-1s* overexpression on innate immune response using multiple methods. First, we used a standard pathogen resistance assay using exposure to *Pseudomonas aeruginosa* (PA14)^47^. Using a canonical PA14 fast kill assay, we found that glutamatergic, octopaminergic, and GABAergic *xbp-1s* animals all displayed a significant increase in survival against PA14, with the octopaminergic *xbp-1s* displaying the most significant increase in survival even after 8 hours (**Fig. 4B**). This is consistent with the octopaminergic *xbp-1s* animals having the greatest change in expression of genes associated with immune response and previous reports that indicate a functional role for octopamine signaling in innate immunity in *C. elegans*^34^. In addition, since octopaminergic *xbp-1s* animals are the only condition that display an improved lifespan, it is possible that the improvement in pathogen resistance of octopaminergic *xbp-1s* animals reaches a critical level to also impact longevity.

**Fig. 4.**
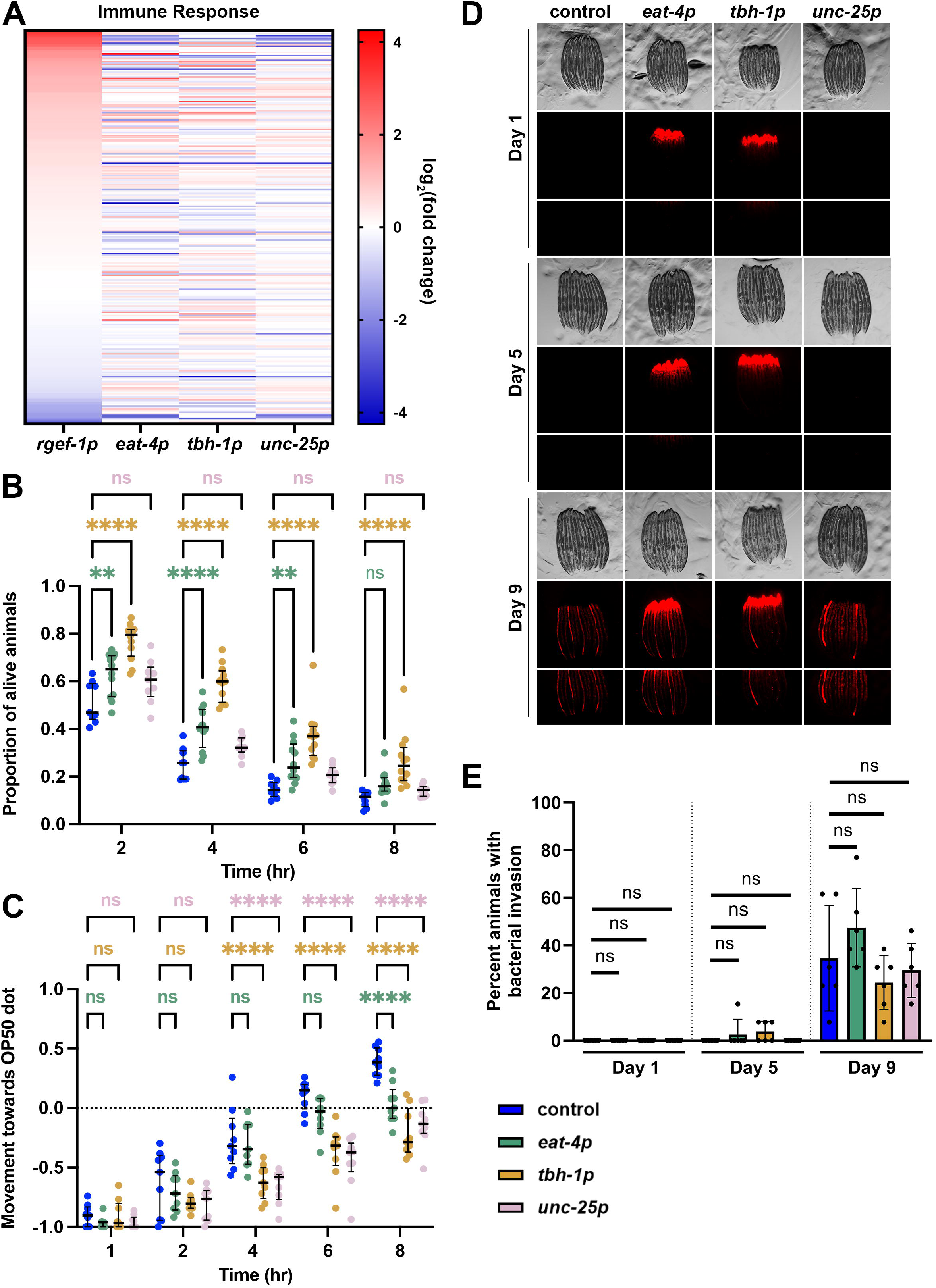
Glutamatergic, octopaminergic, and GABAergic *xbp-1s* enhance pathogen resistance and increases pathogen apathy. **(A)** Heat map of immune response (GO:0006955) gene expression under pan-neuronal (*rgef-1p*), glutamatergic (*eat-4p*), octopaminergic (*tbh-1p*), and GABAergic (*unc-25p*) *xbp-1s* overexpression. Warmer colors indicate increased expression, and cooler colors indicate decreased expression. See **Table S6**. **(B)** Survival analysis of control (N2, blue), glutamatergic *xbp-1s* (green, *eat-4p*), octopaminergic *xbp-1s* (yellow, *tbh-1p*), or GABAergic *xpb-1s* (pink, *unc-25p*) on PA14 fast kill assay plates for 2, 4, 6, and 8 hours. Each fast kill assay is comprised of 3 technical replicates per biological replicate and at least 3 biological replicates per condition. Results were analyzed via two-way ANOVA test; **(p<0.01) ***(p<0.001) ****(p<0.0001). **(C)** Pathogen avoidance behavior of control (N2, blue), glutamatergic *xbp-1s* (green, *eat-4p*), octopaminergic *xbp-1s* (yellow, *tbh-1p*), or GABAergic *xpb-1s* (pink, *unc-25p*) during “forced” food choice assays measured at 1, 2, 3, 6, and 8 hour time points. Each forced food choice assay is comprised of 3 technical replicates per biological replicate and at least 3 biological replicates per condition. Results were analyzed via two-way ANOVA test; **(p<0.01) ***(p<0.001) ****(p<0.0001). **(D)** Representative brightfield and fluorescent images of adult worms grown on bacteria expressing mCherry. Animals are moved to OP50 plates for two hours to remove mCherry expressing bacteria from the intestine before imaging. Any remaining mCherry signal after OP50 clarification are signs of bacterial colonization. **(E)** Quantification of the percent of animals displaying intestinal bacterial colonization was performed across 2 technical replicates for each of 3 biological replicates for a total of 6 replicates. Lines represent mean and standard deviation. * = p ≤ 0.05, ** = p ≤ 0.01, ns = p > 0.05 using a Mann-Whitney test.

*C. elegans* also utilize their nervous system for aversive learning behavior to avoid pathogenic bacteria^48^. This avoidance behavior is mediated by several neurotransmitters, including serotonin^49^ and octopamine^50^, and certain strains with heightened stress responses have been shown to lack this typical avoidance behavior^29^. Here, we used a previously validated forced exposure method^29^ to determine the impact of neuronal *xbp-1s* overexpression on pathogen apathy. Similar to pathogen resistance, glutamatergic, octopaminergic, and GABAergic *xbp-1s* animals all displayed increased apathy to pathogens, with glutamatergic animals having the mildest phenotype (**Fig. 4C**). Thus, it is likely that the heightened resistance to pathogens is directly correlated with a lack of urgency to escape these pathogens. While we did observe an increase in expression of innate immune response genes, it is also possible that the increase in pathogen resistance is due to an increase in gut barrier integrity, as age-associated loss of gut barrier integrity results in infiltration of bacteria and bacterial colonization in the gut^51,52^.

Interestingly, glutamatergic, octopaminergic, and GABAergic *xbp-1s* animals all showed similar breakdown of gut barrier integrity and age-associated bacterial colonization in the gut compared to wild-type controls (**Fig. 4D-E**). These data suggest that the pathogen resistance and apathy of glutamatergic, octopaminergic, and GABAergic *xbp-1s* animals is likely due to a heightened immune response, rather than a gut-barrier-related phenotype.

### *Glutamatergic, octopaminergic, and GABAergic xbp-1s* display reduced lipid levels, but have minor changes to ER morphology

Next, we sought to determine whether neuron subtype-specific *xbp-1s* overexpression altered lipid levels^5,6^. Indeed, we found that *xbp-1s* overexpression resulted in changes to lipid related gene expression (**Fig. 5A**). Therefore, we measured lipid levels using DHS-3::GFP an abundant protein on the surface of *C. elegans* intestinal lipid droplets^53,54^. Interestingly, we saw a significant decrease in lipid droplet abundance in glutamatergic, octopaminergic, or GABAergic *xbp-1s* animals, despite no major changes in lipid droplet size or morphology (**Fig. 5B**). To further evaluate changes in lipid content, we utilized a more comprehensive dye, Oil Red O (ORO), which stains neutral lipids, cholesteryl esters, and lipoproteins^55^. Consistent with lipid droplet imaging, we observed a significant decrease in lipid content in glutamatergic, octopaminergic, or GABAergic *xbp-1s* animals using ORO (**Fig. 5C**). These data suggest that similar to other paradigms of neuronal *xbp-1s* overexpression^3,5,6^, glutamatergic, octopaminergic, or GABAergic *xbp-1s* animals results in depletion of neutral lipids, likely resulting in improved lipid homeostasis.

**Fig. 5.**
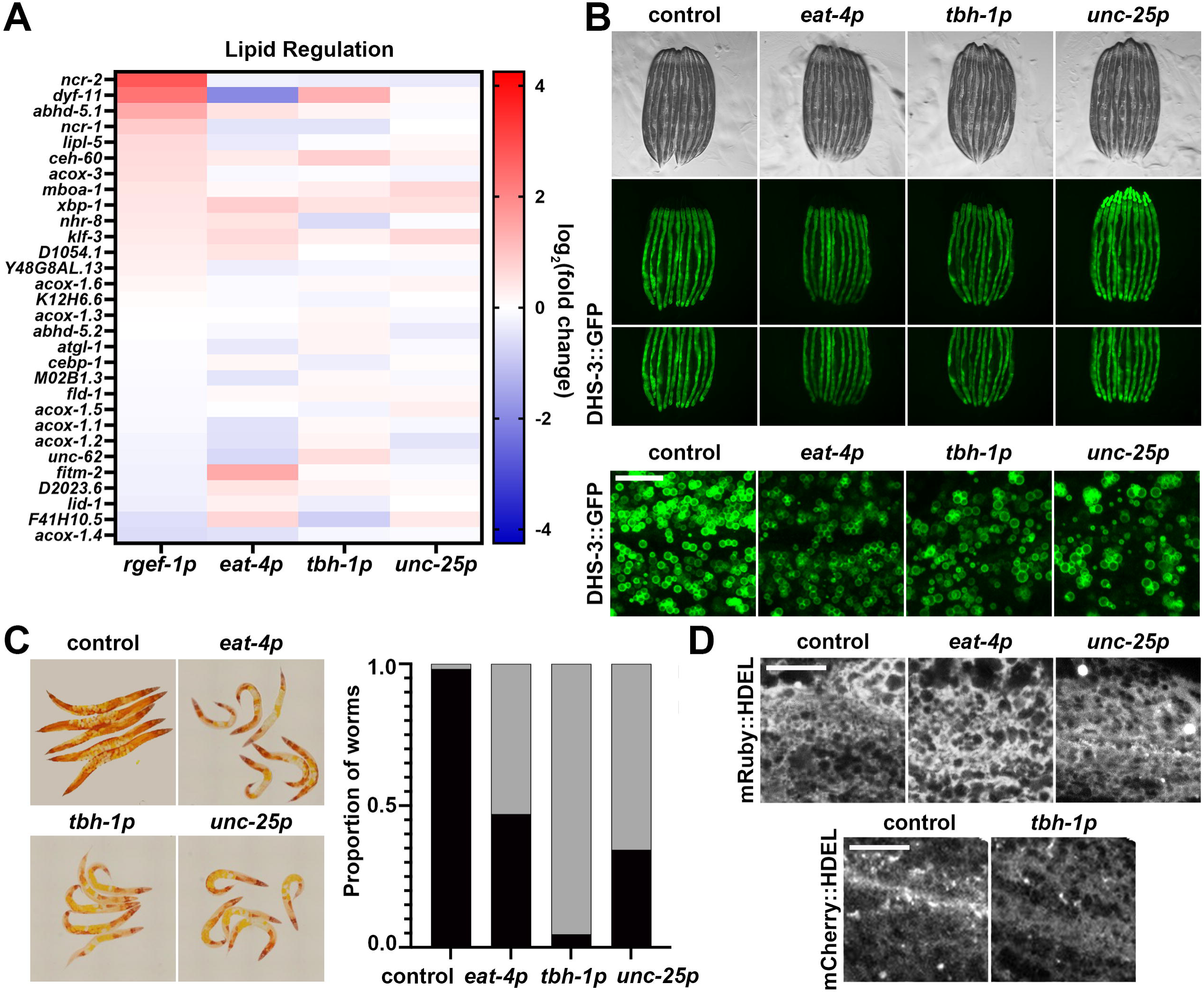
Glutamatergic, octopaminergic, and GABAergic *xbp-1s* results in depletion of lipids. **(A)** Heat map of lipid homeostasis (GO:0055088) gene expression under pan-neuronal (*rgef-1p*), glutamatergic (*eat-4p*), octopaminergic (*tbh-1p*), and GABAergic (*unc-25p*) *xbp-1s* overexpression. Warmer colors indicate increased expression, and cooler colors indicate decreased expression. See **Table S6**. **(B)** Representative fluorescent micrographs of day 3 adult animals of control, glutamatergic *xbp-1s* (*eat-4p*), octopaminergic *xbp-1s* (*tbh-1p*), and GABAergic *xpb-1s* (*unc-25p*) taken on a stereomicroscope and on a confocal microscope (bottom). All images are contrast matched. Scale bar represents 10 um. **(C)** Representative images of day 3 adult animals of control, glutamatergic *xbp-1s* (*eat-4p*), octopaminergic *xbp-1s* (*tbh-1p*), and GABAergic *xpb-1s* (*unc-25p*) of ORO-stained lipids. Quantification of lipid staining as non-lipid depletion (black) and lipid depletion (gray). **(D)** Representative fluorescent micrographs of day 3 adult mRuby::HDEL of control, glutamatergic *xbp-1s* (*eat-4p*), and GABAergic *xbp-1s* (*unc-25p*), animals (top) or day 3 adult mCherry::HDEL of control and octopaminergic *xbp-1s* (*tbh-1p*), animals (bottom). Images are representative of three independent biological replicates and are independently contrast enhanced for each individual image. Scale bar represents 10 µm.

Previous studies have shown that neuronal *xbp-1s* animals exhibit changes in ER morphology associated with a general increase in secretory capacity of the ER and depletion of lipids, potentially through an increase in lipophagy^6^. Therefore, we next sought to determine whether changes in lipid levels found in glutamatergic, octopaminergic, and GABAergic *xbp-1s* animals are also correlated with changes to ER morphology and secretory capacity. To measure general changes to the ER, we first performed imaging of the ER using an mRuby::HDEL fused to an HSP-4 signal sequence to localize the fluorophore to the ER^6^. Since we could not successfully make homozygous octopaminergic *xbp-1s* animals with this mRuby::HDEL marker, we used an mCherry::HDEL fused to a SEL-1 signal sequence, which previous studies have shown display similar ER morphology^6^. Using these ER-localized fluorophores, we did not observe major changes to ER morphology in glutamatergic, octopaminergic, or GABAergic *xbp-1s* animals (**Fig. 5D**). Next, to measure ER secretory capacity, we utilized the yolk protein marker VIT-2::GFP. This maternal yolk protein is secreted by the intestinal ER in adults and subsequently endocytosed by developing eggs and is a commonly used marker for secretory capacity^56,57^. Although fluorescent levels appear to be higher in intact animals (**Fig. S4A**), when we quantitatively measured fluorescent levels in isolated eggs, there was no significant change in VIT-2::GFP signal (**Fig. S4B-C**), suggesting that there are no changes to ER secretory capacity in glutamatergic, octopaminergic, or GABAergic *xbp-1s* animals.

### Glutamatergic xbp-1s promotes protein homeostasis and ER stress resilience

Finally, we measured the impact neuronal subtype *xbp-1s* on protein homeostasis, as UPR^ER^ activation is also directly linked to improved protein homeostasis^7^. We crossed glutamatergic, octopaminergic, and GABAergic overexpressing *xbp-1s* strains into animals expressing fluorescently-tagged aggregation-prone polyglutamine repeats in the intestine^58^ and assessed the extent of aggregation as these animals aged. Strikingly, a significant decrease in fluorescence intensity, demonstrating a reduction of polyQ40 aggregation, was observed at days 1 and 5 of adulthood in all neuronal subtype *xbp-1s* animals as compared to controls (**Fig 6A-B**). Moreover, this was not due to artifacts in our transgenic animal synthesis, as independently synthesized transgenic lines recapitulated these phenotypes (**Fig. 6C-D**). These data suggest that while only glutamatergic *xbp-1s* improved ER proteotoxic stress resistance, all neuronal subtype *xbp-1s* animals have improved peripheral protein homeostasis.

**Fig. 6.**
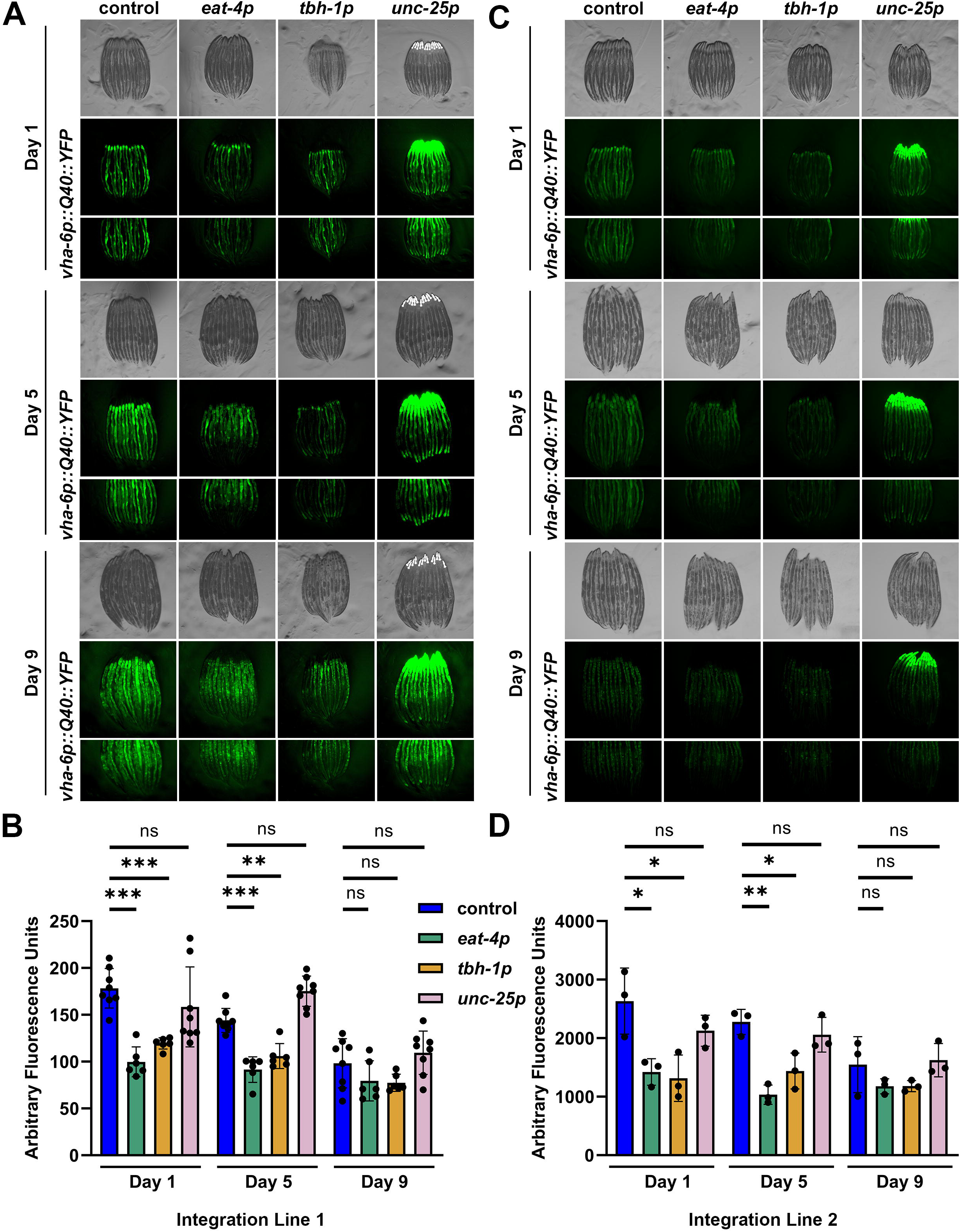
Glutamatergic, octopaminergic, and GABAergic *xbp-1s* enhance proteostasis. **(A)** Representative images of protein aggregation in animals expressing intestinal polyglutamine repeats (*vha-6p::polyQ40::YFP*) ^32^ in glutamatergic (*eat-4p*), octopaminergic (*tbh-1p*), or GABAergic (*unc-25p*) *xbp-1s* animals. All animals were imaged on day 1, 5, and 9 of adulthood. Images were captured using a Leica M205 stereo microscope. **(B)** Quantification of fluorescence integrated density normalized to area was performed across 2 technical replicates each of 3 biological replicates for a total of 6 replicates. Lines represent mean and standard deviation. * = p ≤ 0.05, ** = p ≤ 0.01, ns = p > 0.05 using a Mann-Whitney test. **(C)** Representative images of a second distinct integration line of glutamatergic, octopaminergic, or GABAergic *xbp-1s* animals expressing intestinal polyglutamine repeats (*vha-6p::polyQ40::YFP*) grown on EV or *xbp-1s* RNAi and imaged as per (A). **(D)** Quantification of fluorescence integrated density normalized to area was performed across 3 biological replicates. A Shapiro- Wilk test was used to determine normality and a student’s t-test was used to assess significance.

To investigate the physiological impact of the improved protein homeostasis in these animals, we next measured the impact of neuronal subtype *xbp-1s* on animal survival upon polyQ40 overexpression. Interestingly, octopaminergic *xbp-1s* animals do display a significant increase in lifespan in animals overexpressing polyQ40 (**Fig. S5A**). Similar to standard lifespan extension, this resistance to polyQ overexpression was specific to “normal” sized octopaminergic *xbp-1s* animals, as stunted growth octopaminergic *xbp-1s* animals had reduced lifespan. In addition, glutamatergic and GABAergic *xbp-1s* animals also displayed reduced lifespan in polyQ overexpression conditions. While these data may suggest that octopaminergic *xbp-1s* is the only condition that protects against polyQ overexpression, we found that intestinal polyQ40 overexpression did not consistently reduce lifespan in wild-type animals (**Fig. S5B**). Therefore, it is entirely possible that polyQ40 lifespans do not directly correlate with protein homeostasis capacity.

Therefore, we next measured the ability of neuronal subtype *xbp-1s* animals to resist against protein misfolding stress in the ER. Tunicamycin is a well-characterized ER stressor which blocks N-linked glycosylation in the ER^59^, and animals with neuronal *xbp-1s* exhibit an increased resistance to tunicamycin^2^. Consistent with our transcriptomics data, glutamatergic *xbp-1s* resulted in a small but significant increase in tunicamycin resistance (**Fig. S5C**). Interestingly, octopaminergic and GABAergic *xbp-1s* had no effect on resistance to ER stress (**Fig. S6C-D**), suggesting that even though polyQ aggregation load is decreased in these animals, they do not display a significant impact on proteotoxic stress resilience.

## Conclusions

The UPR^ER^ is involved in diverse cellular processes that impact organismal health, including proteostasis^2,60^, autophagy^7^, lipid metabolism^5,6^, and immune response^61^. Many of these functions can occur in a non-autonomous fashion, whereby neural cells with XBP-1s activation signal to the body to coordinate a homeostatic response^46^. Numerous neural circuits have been implicated in this response, including serotonin, dopamine^3^, tyramine^4^, RIM/RIC interneurons^11^, and glial cells^8^. This study further elucidates this complex neural circuitry, identifying additional functional roles for glutamatergic, octopaminergic, and GABAergic neurons in non-autonomous UPR^ER^ signaling.

The overexpression of *xbp-1s* in glutamatergic, octopaminergic, and GABAergic neurons requires several considerations. First, promoter strength and neuron number can drive different phenotypes across each neuronal *xbp-1s* overexpression paradigm, which is unrelated to the biological significance of a particular neuronal identity. This is especially important to consider as using two different neuron-specific promoters, *rab-3p* and *rgef-1p*, display dramatically different downstream transcriptional responses. However, our data show that *xbp-1s* expression level alone does not purely drive phenotypic outcome. While previous studies utilizing pan-neuronal, serotonergic, or dopaminergic *xbp-1s* showed a significant increase in *xbp-1s* overexpression^3^, in our study, although we can see a trend for an increase in whole-body *xbp- 1s* expression in glutamatergic, octopaminergic, and GABAergic neuron-driven *xbp-1s* overexpression, these data did not reach statistical significance. Despite this lack of a significant increase in *xbp-1s* levels, we still observed dramatic transcriptomic changes, especially in glutamatergic and octopaminergic *xbp-1s*, with changes in expression in canonical XBP-1s targets^62^, protein homeostasis^2^, and immune response^41^ pathways. Further, while there are 79 glutamatergic and 34 GABAergic neurons and glutamate and GABA are the most prevalent excitatory and inhibitory neurotransmitters, respectively, here we only find increased longevity when *xbp-1s* is expressed in a pair of octopaminergic neurons. This finding is consistent with other analogous non-autonomous signaling pathways that have system-wide effects dependent on just a handful of neurons. For example, longevity conferred by dietary restriction, one of the best conserved lifespan-extending paradigms across diverse animal species, is dependent on the two ASI neurons in *C. elegans*^63^. Similarly, non-autonomous *xbp-1s* signaling has been shown to extend lifespan when expressed in a relatively small numbers of cells, including: the four cephalic sheathe glia^8^, the four neurons expressing the tyramine synthesis gene^4^, the six serotonergic neurons, the eight dopaminergic neurons^3^, and in this work, the two octopaminergic neurons. A possible explanation for this is that: 1) it would be inefficient for numerous neurons to redundantly carry out the same function and 2) in order to enact broad effects on metabolism, proteostasis, etc., a sufficiently specific signal is preferred to one that might interfere with the diverse signaling performed by the many glutamatergic and GABAergic neurons. Thus, if we were to overexpress *xbp-1s* in specific subtypes of glutamatergic or GABAergic neurons, we may observe more robust phenotypic effects. Taken together, these data argue that neuron number and promoter strength alone do not drive phenotypic outcomes, and neuronal identity is a critical factor in non-autonomous signaling, even when using an artificial system such as *xbp-1s* overexpression.

Moreover, serotonergic and dopaminergic *xbp-1s*, both of which have lifespan extensions, were shown to be beneficial by eliciting distinct responses in the periphery. In contrast to dopaminergic *xbp-1s* driving lipid depletion, serotonergic *xbp-1s* resulted in an increase in lipids. These two neuronal subtypes, at least in terms of lipid remodeling, have opposing effects.

Beyond neural *xbp-1s*, a similar phenomenon is observed in whole-body *xbp-1s* overexpression wherein no extension in lifespan occurs, which may be due to the cumulative effects of muscle *xbp-1s* shortening lifespan while intestinal and neuronal *xbp-1s* extend lifespan. Altogether, these data suggest that overexpression of *xbp-1s* in a smaller subset of cell types may be more effective at revealing specific downstream pathways, avoiding pleiotropic effects which can occur in more broad *xbp-*1s overexpression.

In a previous study, it was found that overexpressing *xbp-1s* driven by the tyramine synthesis gene, *tdc-1*, was sufficient to extend longevity and drive non-autonomous UPR^ER4^. Two pairs of neurons express tyramine as tyramine is the chemical precursor of octopamine: the octopaminergic RIC neurons and the tyraminergic RIM neurons. It was determined that UPR^ER^ was upregulated in a tyraminergic signaling specific manner. Whether lifespan extension depended on tyramine, octopamine, or a combination of these, however, was left ambiguous as the *tdc-1* promoter used to express *xbp-1s* targets both RIM and RIC neurons. In this study, a potential resolution to this ambiguity is presented. Here, we overexpress *xbp-1s* exclusively in the RIC neurons as opposed to the RIM neurons which signal through both tyramine and glutamate. Thus, our glutamatergic neuron overexpression of *xbp-1s* also targets the tyraminergic RIM neurons, to the exclusion of octopaminergic RIC neurons^33,37^. Here, we find that glutamatergic overexpression of *xbp-1s* increases resilience to tunicamycin, a phenotype related to UPR^ER^. Thus, our findings on glutamatergic *xbp-1s* may be phenotypes driven by the tyraminergic RIM neurons, rather than other glutamatergic neuron subtypes. Further supporting this idea, previous work using a transcriptional reporter in a pan-neuronal *xbp-1s* overexpression animal has shown the tyramine synthesis gene, but not the glutamatergic *vglut* gene, was required for upregulation of UPR^ER4^. As this condition leaves tyraminergic signaling, but not glutamatergic signaling, of RIM neurons intact, it suggests that RIM neurons drive UPR^ER^ in a tyramine-dependent manner, but not lifespan, which results from octopaminergic signaling by RIC neurons.

Interestingly, we find that although GABAergic *xbp-1s* had minimal changes to gene expression, these animals still displayed significant changes to organismal health, including improved proteostasis and immune response. While transcriptional changes are not the only change that could translate to physiology, as altered protein function, organelle dynamics, and metabolism can all occur in the absence of transcriptional change, technical limitations could also be responsible for the lack of difference observed in GABAergic *xbp-1s*. Here, we used whole-worm transcriptomics, and it is entirely possible that opposite changes in gene expression in different tissues could result in a net result of no change. Indeed, in terms of *xbp-1s* overexpression, this is observed wherein whole-body overexpression of *xbp-1s* does not result in lifespan extension, likely due to the summation of negative effects in the muscle and positive effects in the intestine and neurons^2^. Thus, it is entirely possible that GABAergic *xbp-1s* may drive differential effects in different tissue, as it does drive depletion of lipids and increased protein homeostasis in the intestine. Future tissue-specific studies can reveal whether these physiological outputs are dependent on gene expression changes in the intestine in these animals.

A potential limitation of our findings is whether ectopic gene overexpression correlates with the endogenous roles these neuronal subtypes play in signaling or gene regulation. Previous work in numerous animal models provide sufficient evidence that this is the case. In *C. elegans*, olfactory sensation of pathogenic bacteria utilize neuron-to-body XBP-1s signaling through TGF- β signaling to improve longevity and healthspan^11^. In mice, Xbp1s overexpression in POMC neurons promotes adipose tissue UPR^ER^ to improve metabolic health^9^. Similarly, hepatic *Xbp1s* activation in mice promotes metabolic health downstream of food perception^10^. In *D. melanogaster*, glutamate signaling can promote lipid mobilization as a systemic metabolite, altering lipid metabolism^64^. These reports suggest that even ectopic genetic models can provide mechanistic insight into important endogenous physiological processes.

Lastly, in addition to neuronal identity, there exist complex neuronal circuits that involve signaling between different neuronal subtypes – some of which are discussed here – which may contribute to phenotypic differences. In this study, we are generalizing neuronal signals as separate entities, although neuronal circuits are often intertwined and complex ^65^. While our previous study has separated the utility of dopamine in serotonergic *xbp-1s* signaling and vice versa ^3^, this does not preclude the convergence of other neuronal subtypes. For example, we find that although glutamatergic *xbp-1s* animals display mostly changes to protein homeostasis-related pathways, they also show improved immune function and increased lipid depletion.

While the improvement in protein homeostasis could be responsible for increased immune function ^41^, it is entirely possible that glutamatergic neurons can also recruit octopaminergic or GABAergic signaling to alter immune response and lipid metabolism. Future studies mapping the neural circuitry across subtypes will be necessary to develop a full neural map of non-autonomous XBP-1s signaling. Overall, our study further builds on the complex literature of non-autonomous XBP-1s signaling, adding three additional neuronal subtypes to this rapidly expanding map.

## Acknowledgements

A.J.C., A.J.H., M.A., and G.G. are supported by T32AG052374; A.J.H. is supported by the NSF GRFP; S.P.C. is supported by NIH R01AG058610 and Hevolution Foundation award 748 HF AGE-004; B.A.B. is supported by the Simons Collaboration on Plasticity in the Aging Brain grant SF811217; P.J.M. is supported by the Impetus Grants 014746-00001 and the Baxter award; and R.H.S. is supported by R00AG065200 and R01AG079806 from the National Institute on Aging and the Glenn Foundation for Medical Research and AFAR Grant for Junior Faculty Award. Some strains were provided by the CGC, which is funded by the NIH Office of Research Infrastructure Programs (P40 OD010440). Some gene analysis was performed using Wormbase, which is funded on a U41 grant HG002223.

## Competing Financial Interests

All authors of the manuscript declare that they have no competing interests.

## Data Availability

All data required to evaluate the conclusions in this manuscript are available within the manuscript and Supporting Information. All strains synthesized in this manuscript are derivatives of N2 or other strains from CGC and are either made available on CGC or available upon request. All raw RNA-seq datasets are available through Annotare 2.0 Array Express Accession E-MTAB-14132.

---

**Fig. S1.**
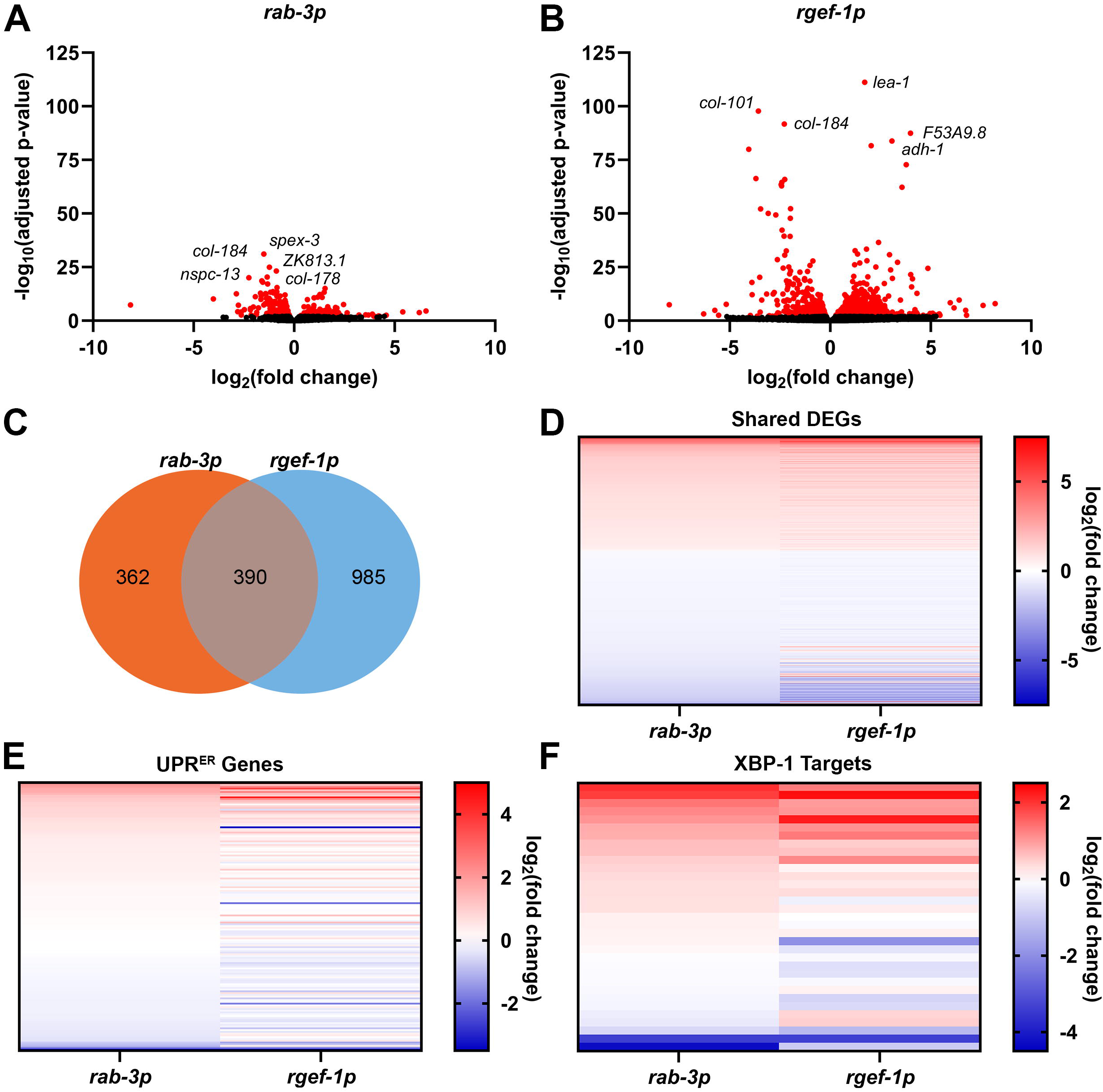
A comparison of pan-neuronal *xbp-1s* expression via two different promoters reveals differences in whole-body transcriptomic changes. Volcano plots of whole-body genome-wide changes in gene expression upon pan-neuronal *xbp-1s* overexpression driven by (A) *rab-3* promoter or (B) *rgef-1* promoter. Red dots indicate significantly differentially expressed genes with p-value ≤ 0.01. See **Table S3** for a list of differentially expressed genes and expression values. **(C)** Comparison of differentially expressed genes (p-value ≤ 0.01) between worms expressing *xbp-1s* pan-neuronally driven by *rab-3p* or *rgef-1p*. For a complete list of differentially expressed genes in each group, see **Table S4**. **(D)** Heat map of common differentially expressed genes upon pan-neuronal *xbp-1s* expression under control of *rab-3* or *rgef-1* promoter. Warmer colors indicate increased expression, and cooler colors indicate decreased expression. See **Table S5** for a list of genes and values. **(E)** Heat map of UPR^ER^ related gene (GO:0030968) expression under pan-neuronal *xbp-1s* driven by *rab-3* or *rgef-1* promoter. Warmer colors indicate increased expression, and cooler colors indicate decreased expression. See **Table S5** for a list of genes and values. **(F)** Heat map of XBP-1s target gene ^31^ expression under pan-neuronal *xbp-1s* driven by *rab-3* or *rgef-1* promoter. Warmer colors indicate increased expression, and cooler colors indicate decreased expression. See **Table S5** for a list of genes and values.

**Figure S2.**
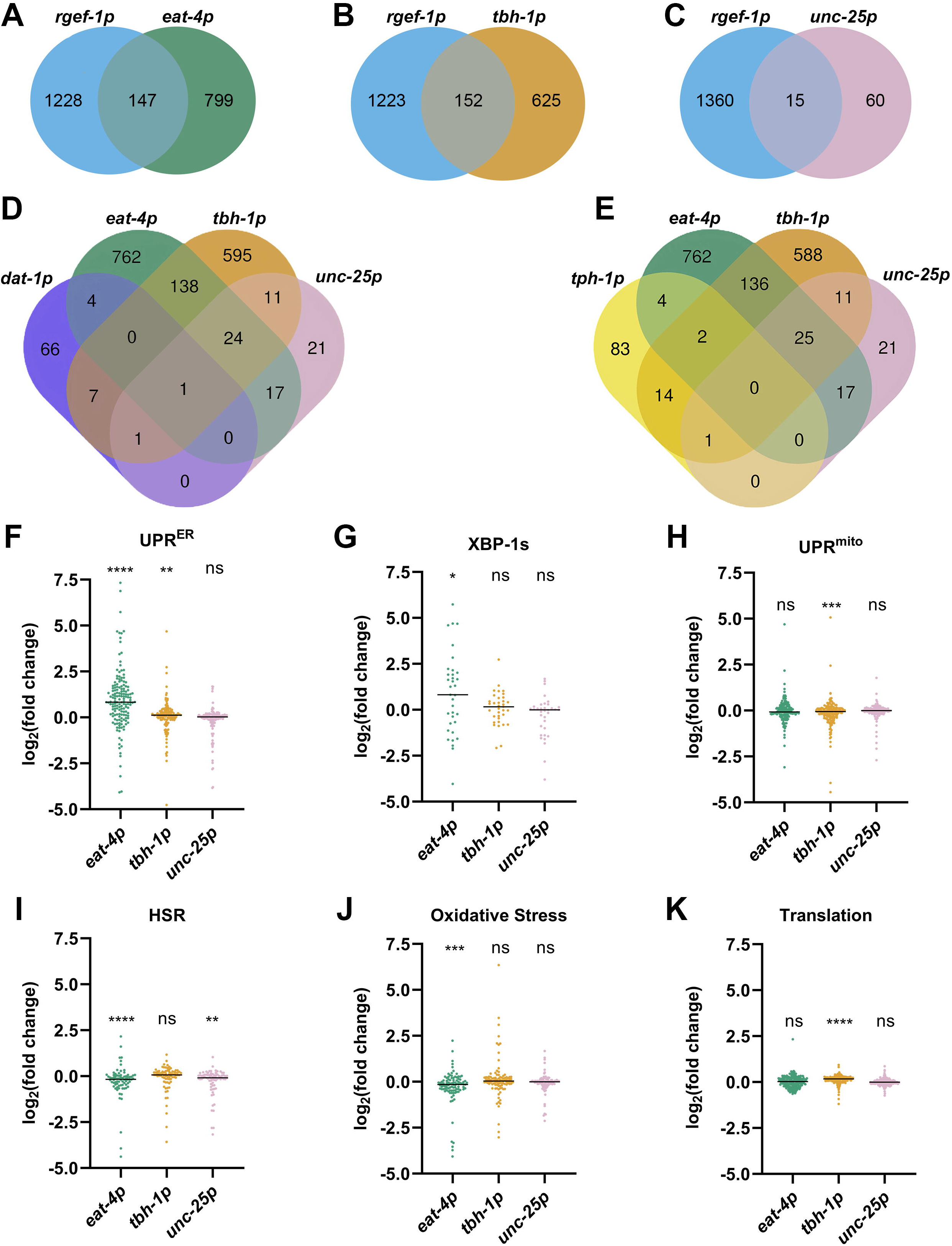
*xbp-1s* overexpression in *C. elegans* glutamatergic, octopaminergic, and GABAergic neurons drives differential gene expression. Comparison of differentially expressed genes (p-value ≤ 0.01) in neuronal *xbp-1s* driven by *rgef-1p* and **(A)** glutamatergic (*eat-4p*), **(B)** octopaminergic *(tbh-1p*), and **(C)** GABAergic (*unc-25p*) *xbp-1s* expression. For a complete list of differentially expressed genes in each group, see **Table S3**. Comparison of differentially expressed genes (p-value ≤ 0.01) in glutamatergic, octopaminergic, GABAergic, and **(D)** dopaminergic or **(E)** serotonergic neurons. See **Table S4**. Gene expression changes in groups of genes related to **(F)** UPR^ER^ (GO:0030968), **(G)** XBP-1s targets ^31^, **(H)** mitochondrial unfolded protein response (GO:0034514), **(I)** heat shock response (GO:0009408), **(J)** oxidative stress response (GO:0006979), and **(K)** translation (GO:0006412). Significance was determined using a one-sample Wilcoxon test. * = p ≤ 0.05, ** = p ≤ 0.01, *** = p ≤ 0.001, **** = p ≤ 0.0001, ns = p > 0.05

**Fig. S3.**
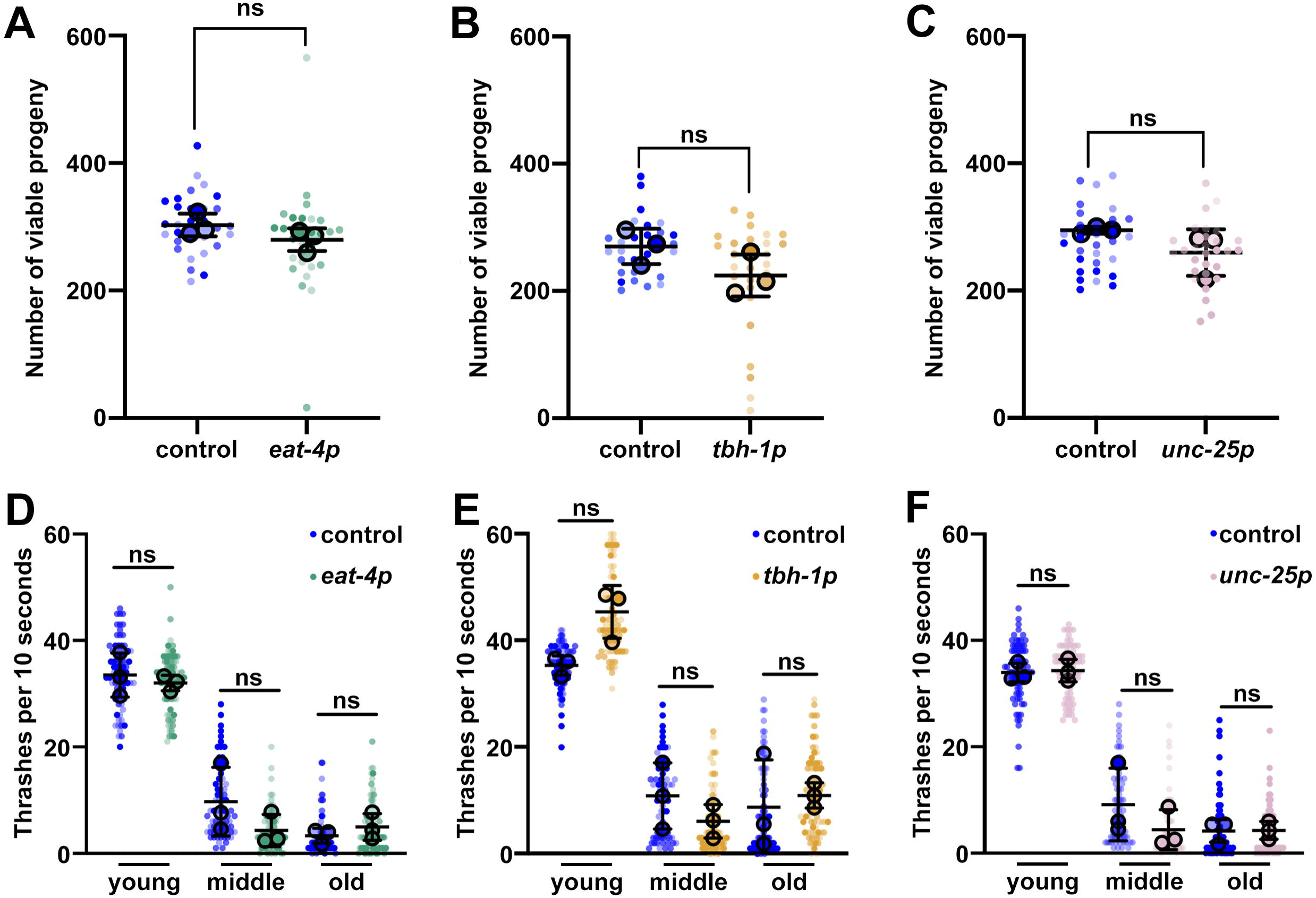
Glutamatergic, octopaminergic, and GABAergic *xbp-1s* does not increase healthspan. Measurements of fecundity of control (blue) and **(A)** glutamatergic *xbp-1s* (green, *eat-4p*), **(B)** octopaminergic *xbp-1s* (yellow, *tbh-1p*), and **(C)** GABAergic *xbp-1s* (pink, *unc-25p*) animals. Total number of eggs that hatched were counted per animal. Measurements of thrashing of control (blue) and **(D)** glutamatergic *xbp-1s* (green, *eat-4p*), **(E)** octopaminergic *xbp-1s* (yellow, *tbh-1p*), and **(F)** GABAergic *xbp-1s* (pink, *unc-25p*) animals. Number of thrashes was assessed over a 10 second period in animals at day 1 adult (young), day 4-5 adult (middle), and day 9 adult (old) in M9 solution with each thrash being counted as a movement from a concave to a convex formation. For SuperPlots, each small dot represents a single animal with various intensities of colors representing independent biological replicates and each large dot is the median value of each biological replicate. Lines represent the median across all biological replicates and whiskers indicate interquartile range. Statistical analysis was performed using a Mann-Whitney test.

**Fig. S4.**
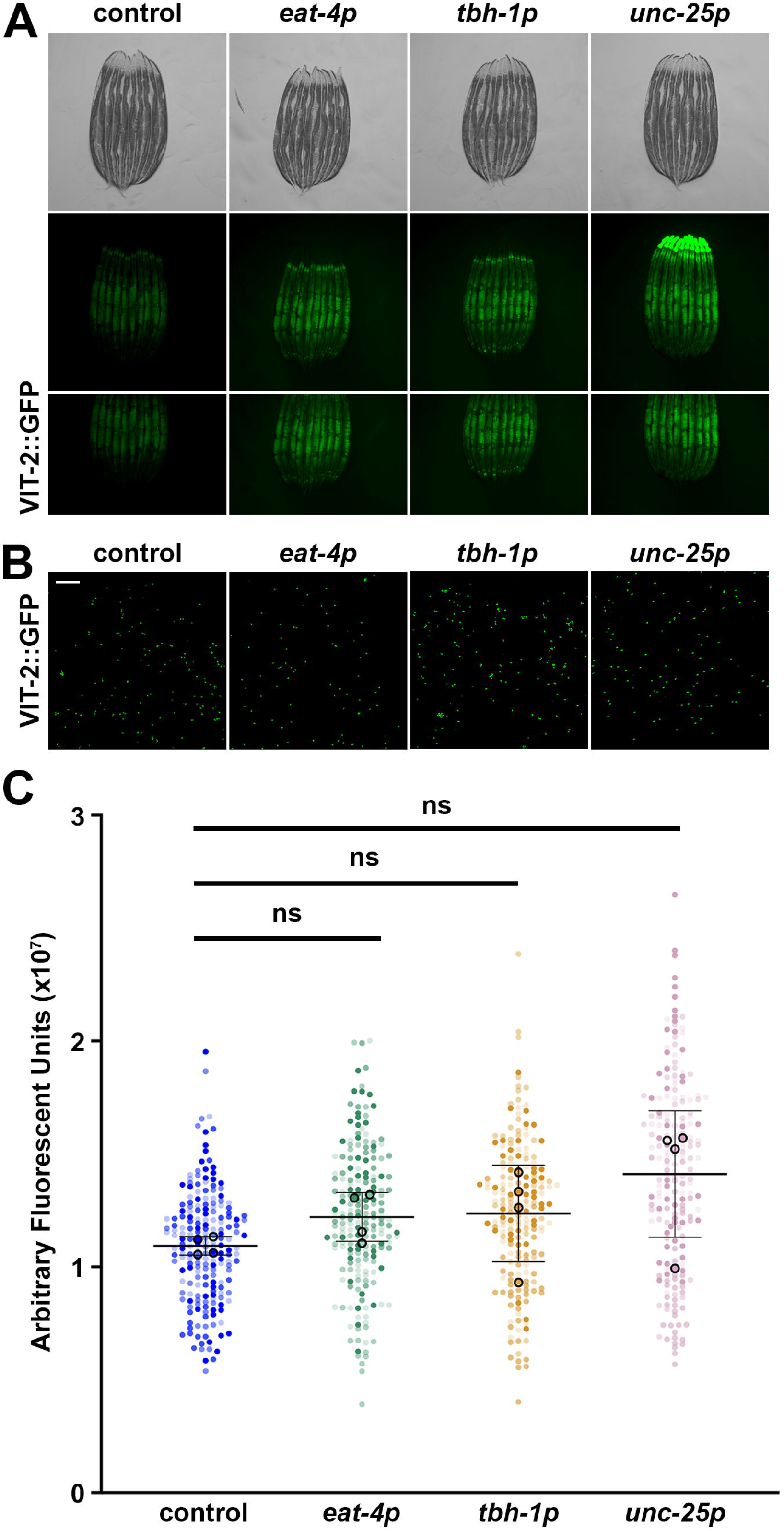
Glutamatergic, octopaminergic, and GABAergic *xbp-1s* does not alter ER secretory capacity. **(A)** Representative fluorescent micrographs of day 3 adult animals of VIT2::GFP in control, glutamatergic *xbp-1s* (*eat-4p*), octopaminergic *xbp-1s* (*tbh-1p*), or GABAergic *xpb-1s* (*unc-25p*). All images are contrast-matched. **(B)** Representative fluorescent micrographs of eggs collected using a standard bleaching protocol of day 3 adult animals of VIT2::GFP in control, glutamatergic *xbp-1s* (*eat-4p*), octopaminergic *xbp-1s* (*tbh-1p*), or GABAergic *xpb-1s* (*unc-25p*). All images are contrast-matched. Scale bar represents 500 µm.**(C)** Quantification of eggs from (B) using measurements of integrated intensity. For SuperPlots, each small dot represents a single animal with various intensities of colors representing independent biological replicates and each large dot is the median value of each biological replicate. Lines represent the median across all biological replicates and whiskers indicate interquartile range. Statistical analysis was performed using a Mann-Whitney test.

**Fig. S5.**
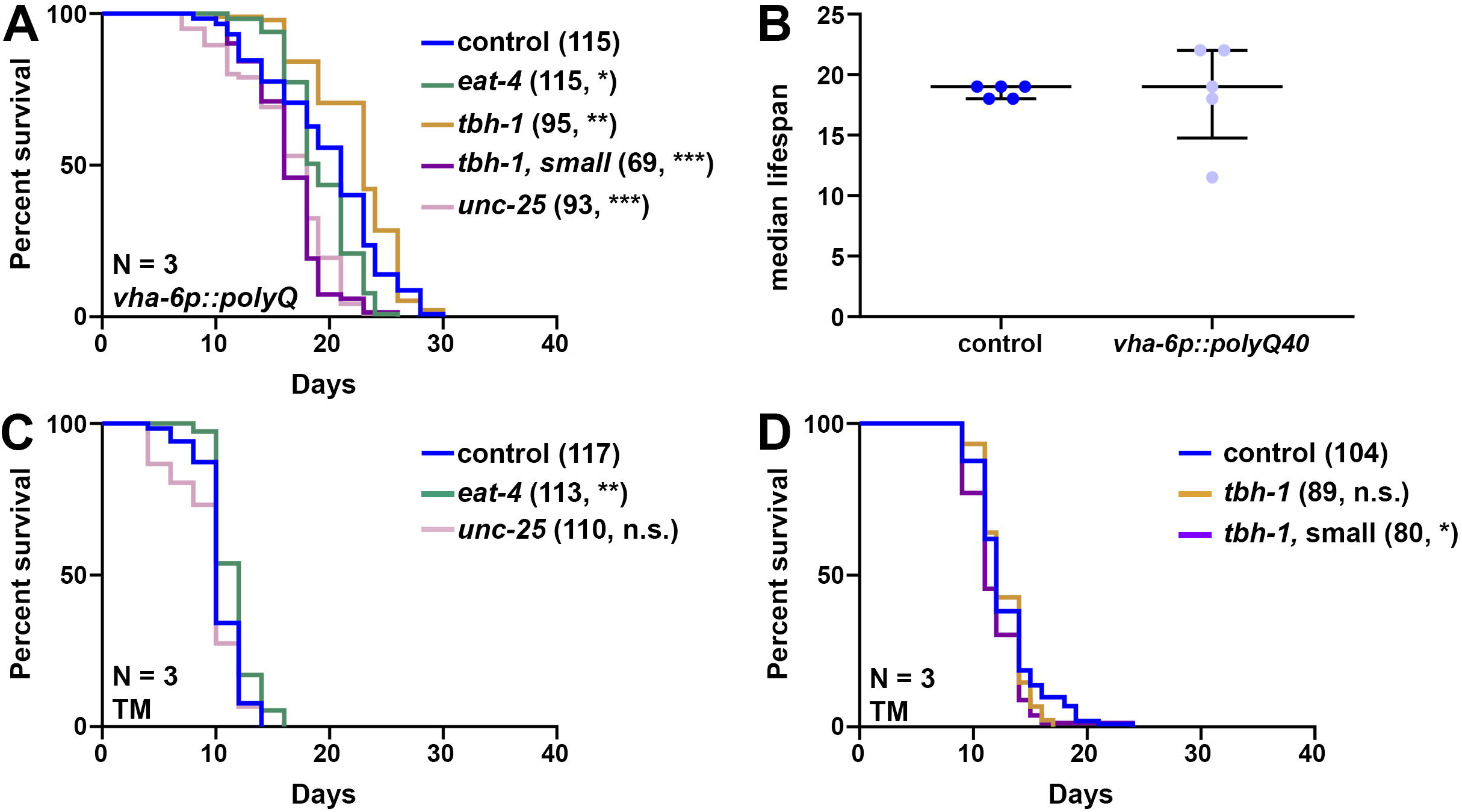
Octopaminergic *xbp-1s* promotes lifespan in polyQ40 expression animals, but not on tunicamycin. **(A)** Lifespan measurements of control (here, *vha-6p::polyQ*), blue), glutamatergic *xbp-1s* animals (*eat-4p*, green), octopaminergic *xbp-1s* separated for normal sized (*tbh-1,* yellow) or stunted growth (*tbh-1,* small, purple), and GABAergic *xbp-1s* (*unc-25p,* pink) animals expressing polyQ40::YFP in the intestine (*vha-6p::polyQ*). **(B)** Median lifespan measurements from 5 replicates of wild-type N2 animals either expressing *vha-6p::polyQ40* (light blue, *vha-6p::polyQ*) or not (dark blue, control). Dots indicate each biological replicate and lines represent median plus interquartile range. **(C)** Lifespan measurements of control (blue, light blue), glutamatergic *xbp-1s* animals (*eat-4p*, green), and GABAergic *xbp-1s* (*unc-25p,* pink) animals moved onto 25 mg/mL tunicamycin (TM) plates starting from day 1 of adulthood. Lifespans were scored every 2 days and data is representative of 3 biological replicates (N). **(D)** Lifespan measurements of control (blue) and octopaminergic *xbp-1s* animals moved onto 25 mg/mL tunicamycin (TM) plates starting from day 1 of adulthood. Octopaminergic *xbp-1s* animals were separated into normal size (yellow, *tbh-1p*) and stunted growth (purple, *tbh-1p,* small). Sample size (n) is written next to each condition followed by significance measuredusing Log-Rank testing: n.s. = not significant, * = p < 0.05, *** = p < 0.001. All statistical analysis is available in **Table S7**.

Table S1. Strains used in this study. Table S2. Primers used in this study.

Table S3. List of differentially expressed genes. Table S4. List of all genes in Venn Diagrams.

Table S5. List of all genes used in heat maps. Table S6. List of all gene ontologies.

Table S7. Lifespan statistics.

## Notes

### Competing Interest Statement

The authors have declared no competing interest.

### Summary of Updates

New data has been added that revealed that octopaminergic xbp-1s animals are a bimodal population with a long-lived and short-lived cohort. The entire manuscript was redone using this long-lived cohort to show all the benefits of octopaminergic xbp-1s.

## References

1. Dutta, N., Garcia, G. & Higuchi-Sanabria, R. Hijacking Cellular Stress Responses to Promote Lifespan. *Front*. Aging 3, 860404 (2022).

2. Taylor, R. C. & Dillin, A. XBP-1 Is a Cell-Nonautonomous Regulator of Stress Resistance and Longevity. Cell 153, 1435–1447 (2013).

3. Higuchi-Sanabria, R. et al. Divergent Nodes of Non-autonomous UPRER Signaling through Serotonergic and Dopaminergic Neurons. Cell Reports 33, 108489 (2020).

4. Özbey, N. P. et al. Tyramine Acts Downstream of Neuronal XBP-1s to Coordinate Inter-tissue UPRER Activation and Behavior in C. elegans. Developmental Cell 55, 754–770.e6 (2020).

5. Imanikia, S., Sheng, M., Castro, C., Griffin, J. L. & Taylor, R. C. XBP-1 Remodels Lipid Metabolism to Extend Longevity. Cell Reports 28, 581–589.e4 (2019).

6. Daniele, J. R. et al. UPRER promotes lipophagy independent of chaperones to extend life span. Science Advances 6, eaaz1441 (2020).

7. Imanikia, S., Özbey, N. P., Krueger, C., Casanueva, M. O. & Taylor, R. C. Neuronal XBP-1 Activates Intestinal Lysosomes to Improve Proteostasis in C. elegans. Current Biology 29, 2322–2338.e7 (2019).

8. Frakes, A. E. et al. Four glial cells regulate ER stress resistance and longevity via neuropeptide signaling in C. elegans. Science 367, 436–440 (2020).

9. Williams, K. W. et al. Xbp1s in Pomc Neurons Connects ER Stress with Energy Balance and Glucose Homeostasis. Cell Metabolism 20, 471–482 (2014).

10. Brandt, C. et al. Food Perception Primes Hepatic ER Homeostasis via Melanocortin-Dependent Control of mTOR Activation. Cell 175, 1321–1335.e20 (2018).

11. De-Souza, E. A., Thompson, M. A. & Taylor, R. C. Olfactory chemosensation extends lifespan through TGF-β signaling and UPR activation. Nat Aging 3, 938–947 (2023).

12. Bar-Ziv, R. et al. Measurements of Physiological Stress Responses in C. Elegans. JoVE (Journal of Visualized Experiments) e61001 (2020) doi:10.3791/61001.

13. Garcia, G., Homentcovschi, S., Kelet, N. & Higuchi-Sanabria, R. Imaging of Actin Cytoskeletal Integrity During Aging in C. elegans. in Cytoskeleton : Methods and Protocols (ed. Gavin, R. H.) 101–137 (Springer US, New York, NY, 2022). doi:10.1007/978-1-0716-1661-1_5.

14. Schindelin, J., et al. Fiji: an open-source platform for biological-image analysis. Nat Methods 9, 676–682 (2012).

15. Egge, N. et al. Age-Onset Phosphorylation of a Minor Actin Variant Promotes Intestinal Barrier Dysfunction. Developmental Cell 51, 587–601.e7 (2019).

16. Grant, B. & Hirsh, D. Receptor-mediated Endocytosis in the Caenorhabditis elegans Oocyte. MBoC 10, 4311–4326 (1999).

17. Lord, S., Velle, K., Mullins, D. & Fritz-Laylin, L. SuperPlots: Communicating reproducibility and variability in cell biology. Journal of Cell Biology 219, e202001064 (2020).

18. Stuhr, N. et al. Rapid Lipid Quantification in Caenorhabditis elegans by Oil Red O and Nile Red Staining. BIO-PROTOCOL 12, (2022).

19. Lynn, D. A. et al. Omega-3 and -6 fatty acids allocate somatic and germline lipids to ensure fitness during nutrient and oxidative stress in Caenorhabditis elegans. Proceedings of the National Academy of Sciences 112, 15378–15383 (2015).

20. Dobin, A. et al. STAR: ultrafast universal RNA-seq aligner. Bioinformatics 29, 15–21 (2013).

21. Love, M. I., Huber, W. & Anders, S. Moderated estimation of fold change and dispersion for RNA-seq data with DESeq2. Genome Biol 15, 550 (2014).

22. Calfon, M. et al. IRE1 couples endoplasmic reticulum load to secretory capacity by processing the XBP-1 mRNA. Nature 415, 92–96 (2002).

23. Chen, E. Y. et al. Enrichr: interactive and collaborative HTML5 gene list enrichment analysis tool. BMC Bioinformatics 14, 1–14 (2013).

24. Kuleshov, M. V. et al. Enrichr: a comprehensive gene set enrichment analysis web server 2016 update. Nucleic Acids Research 44, W90–W97 (2016).

25. Stroustrup, N. et al. The Caenorhabditis elegans Lifespan Machine. Nature Methods 10, 665–670 (2013).

26. Torres, T. C. et al. Surveying Low-Cost Methods to Measure Lifespan and Healthspan in Caenorhabditis elegans. JoVE (Journal of Visualized Experiments) e64091 (2022) doi:10.3791/64091.

27. Cezairliyan, B. et al. Identification of Pseudomonas aeruginosa Phenazines that Kill Caenorhabditis elegans. PLOS Pathogens 9, e1003101 (2013).

28. Mahajan-Miklos, S., Tan, M.-W., Rahme, L. G. & Ausubel, F. M. Molecular Mechanisms of Bacterial Virulence Elucidated Using a Pseudomonas aeruginosa– Caenorhabditis elegans Pathogenesis Model. Cell 96, 47–56 (1999).

29. Nair, T., Weathers, B. A., Stuhr, N. L., Nhan, J. D. & Curran, S. P. Serotonin deficiency from constitutive SKN-1 activation drives pathogen apathy. Preprint at 10.1101/2024.02.10.579755 (2024).

30. Stuhr, N. L., Ramos, C. M., Turner, C. D., Soukas, A. A. & Curran, S. P. C. elegans display antipathy behavior towards food after contemporaneous integration of nutritional needs and dietary lipid availability. 2024.02.23.581740 Preprint at 10.1101/2024.02.23.581740 (2024).

31. Urano, F. et al. A survival pathway for Caenorhabditis elegans with a blocked unfolded protein response. Journal of Cell Biology 158, 639–646 (2002).

32. Bar-Ziv, R. et al. Glial-derived mitochondrial signals affect neuronal proteostasis and aging. Science Advances 9, eadi1411 (2023).

33. Serrano-Saiz, E. et al. Modular Control of Glutamatergic Neuronal Identity in C. elegans by Distinct Homeodomain Proteins. Cell 155, 659–673 (2013).

34. Sellegounder, D., Yuan, C.-H., Wibisono, P., Liu, Y. & Sun, J. Octopaminergic Signaling Mediates Neural Regulation of Innate Immunity in Caenorhabditis elegans. mBio 9, e01645–18 (2018).

35. McIntire, S. L., Jorgensen, E. & Horvitz, H. R. Genes required for GABA function in Caenorhabditis elegans. Nature 364, 334–337 (1993).

36. Lee, R. Y. N., Sawin, E. R., Chalfie, M., Horvitz, H. R. & Avery, L. EAT-4, a Homolog of a Mammalian Sodium-Dependent Inorganic Phosphate Cotransporter, Is Necessary for Glutamatergic Neurotransmission in Caenorhabditis elegans. J Neurosci 19, 159–167 (1999).

37. Alkema, M. J., Hunter-Ensor, M., Ringstad, N. & Horvitz, H. R. Tyramine Functions Independently of Octopamine in the Caenorhabditis elegans Nervous System. Neuron 46, 247–260 (2005).

38. Jin, Y., Jorgensen, E., Hartwieg, E. & Horvitz, H. R. The Caenorhabditis elegans Gene unc- 25Encodes Glutamic Acid Decarboxylase and Is Required for Synaptic Transmission But Not Synaptic Development. J Neurosci 19, 539–548 (1999).

39. Gelino, S. et al. Intestinal Autophagy Improves Healthspan and Longevity in C. elegans during Dietary Restriction. PLoS Genet 12, e1006135 (2016).

40. Yang, Y. et al. Autophagy protein ATG-16.2 and its WD40 domain mediate the beneficial effects of inhibiting early-acting autophagy genes in C. elegans neurons. Nat Aging 4, 198– 212 (2024).

41. Richardson, C. E., Kooistra, T. & Kim, D. H. An essential role for XBP-1 in host protection against immune activation in C. elegans. Nature 463, 1092–1095 (2010).

42. Kwon, S., Kim, E. J. E. & Lee, S.-J. V. Mitochondria-mediated defense mechanisms against pathogens in Caenorhabditis elegans. BMB Rep 51, 274–279 (2018).

43. Liu, Y. & Chang, A. Heat shock response relieves ER stress. EMBO J. 27, 1049–1059 (2008).

44. Glover-Cutter, K. M., Lin, S. & Blackwell, T. K. Integration of the Unfolded Protein and Oxidative Stress Responses through SKN-1/Nrf. PLOS Genetics 9, e1003701 (2013).

45. Anderson, N. & Haynes, C. M. Folding the Mitochondrial UPR into the Integrated Stress Response. Trends Cell Biol 30, 428–439 (2020).

46. Homentcovschi, S. & Higuchi-Sanabria, R. A neuron’s ambrosia: non-autonomous unfolded protein response of the endoplasmic reticulum promotes lifespan. Neural Regen Res 17, 309–310 (2022).

47. Tan, M.-W., Mahajan-Miklos, S. & Ausubel, F. M. Killing of Caenorhabditis elegans by Pseudomonas aeruginosa used to model mammalian bacterial pathogenesis. Proceedings of the National Academy of Sciences 96, 715–720 (1999).

48. Meisel, J. D. & Kim, D. H. Behavioral avoidance of pathogenic bacteria by *Caenorhabditis elegans*. Trends in Immunology 35, 465–470 (2014).

49. Zhang, Y., Lu, H. & Bargmann, C. I. Pathogenic bacteria induce aversive olfactory learning in Caenorhabditis elegans. Nature 438, 179–184 (2005).

50. O’Donnell, M. P., Fox, B. W., Chao, P.-H., Schroeder, F. C. & Sengupta, P. A neurotransmitter produced by gut bacteria modulates host sensory behaviour. Nature 583, 415–420 (2020).

51. Egge, N. et al. Age-Onset Phosphorylation of a Minor Actin Variant Promotes Intestinal Barrier Dysfunction. Developmental Cell 51, 587–601.e7 (2019).

52. Garcia, G. et al. Large-scale genetic screens identify BET-1 as a cytoskeleton regulator promoting actin function and life span. Aging Cell 22, e13742 (2023).

53. Na, H. et al. Identification of lipid droplet structure-like/resident proteins in Caenorhabditis elegans. Biochimica et Biophysica Acta (BBA) - Molecular Cell Research 1853, 2481–2491 (2015).

54. Zhang, P. et al. Proteomic study and marker protein identification of Caenorhabditis elegans lipid droplets. Mol. Cell Proteomics 11, 317–328 (2012).

55. Du, J., Zhao, L., Kang, Q., He, Y. & Bi, Y. An optimized method for Oil Red O staining with the salicylic acid ethanol solution. Adipocyte 12, 2179334.

56. Grant, B. & Hirsh, D. Receptor-mediated endocytosis in the Caenorhabditis elegans oocyte. Mol. Biol. Cell 10, 4311–4326 (1999).

57. Stevens, J. & Spang, A. N-glycosylation is required for secretion and mitosis in C. elegans. PLoS ONE 8, e63687 (2013).

58. Morley, J. F., Brignull, H. R., Weyers, J. J. & Morimoto, R. I. The threshold for polyglutamine-expansion protein aggregation and cellular toxicity is dynamic and influenced by aging in Caenorhabditis elegans. Proceedings of the National Academy of Sciences 99, 10417–10422 (2002).

59. Heifetz, A., Keenan, R. W. & Elbein, A. D. Mechanism of action of tunicamycin on the UDP-GlcNAc:dolichyl-phosphate Glc-NAc-1-phosphate transferase. Biochemistry 18, 2186–2192 (1979).

60. Frakes, A. E. & Dillin, A. The UPR(ER): Sensor and Coordinator of Organismal Homeostasis. Mol. Cell 66, 761–771 (2017).

61. Grootjans, J., Kaser, A., Kaufman, R. J. & Blumberg, R. S. The unfolded protein response in immunity and inflammation. Nat Rev Immunol 16, 469–484 (2016).

62. Shen, X., Ellis, R. E., Sakaki, K. & Kaufman, R. J. Genetic interactions due to constitutive and inducible gene regulation mediated by the unfolded protein response in C. elegans. PLoS Genet. 1, e37 (2005).

63. Bishop, N. A. & Guarente, L. Two neurons mediate diet-restriction-induced longevity in C. elegans. Nature 447, 545–549 (2007).

64. Zhao, X. & Karpac, J. Glutamate metabolism directs energetic trade-offs to shape host-pathogen susceptibility in Drosophila. Cell Metabolism 33, 2428–2444.e8 (2021).

65. Randi, F., Sharma, A. K., Dvali, S. & Leifer, A. M. Neural signal propagation atlas of Caenorhabditis elegans. Nature 623, 406–414 (2023).

