## Supplementary material for "Distinct mechanisms of non-autonomous UPR^ER^ mediated by GABAergic, glutamatergic, and octopaminergic neurons": Table S2

| **Strains used in this study** | | |
| --- | --- | --- |
| *C. elegans*: Bristol (N2) strain as wild type (WT) | CGC | N2 |
| *C. elegans*: AGD1395: *uthIs393[vha-6p::Q40::YFP+rol-6(su1006)]* | Dillin Lab, [66] |  |
| *C. elegans*: AGD1415: *pwIs23[vit-2::GFP]* | Dillin lab, [6] |  |
| *C. elegans*: AGD2192: *unc-119(ed3) III; uthSi60[vha-6p::ERss::mRuby::HDEL::unc-54 3'UTR cb-unc-119(+)] IV* | Dillin lab, [6] |  |
| *C. elegans*: MAH602: *sqIs61[vha-6p::Q44::YFP + rol-6(su1006)];* | Hansen Lab, [31] |  |
| *C. elegans*: RHS10: *ldrIs[dhs-3p::dhs-3::GFP + unc-76(+)]* | This study |  |
| *C. elegans*: RHS17: *sybIs3954[tbh-1p::xbp-1s, myo-2p::mCherry]* | This study |  |
| *C. elegans*: RHS18: *sybIs3923[eat-4p::xbp-1s, myo-2p::mCherry]* | This study |  |
| *C. elegans*: RHS19: *glp-4(bn2) I* | This study | SS104 6x backcross |
| *C. elegans*: RHS20: *sybIs3954[tbh-1p::xbp-1s, myo-2p::mCherry], ldrIs[dhs-3p::dhs-3::GFP + unc-76(+)]* | This study |  |
| *C. elegans*: RHS21: *sybIs3954[tbh-1p::xbp-1s, myo-2p::mCherry], pwIs23[vit-2::GFP]* | This study |  |
| *C. elegans*: RHS57: *sybIs3970[unc-25p::xbp-1s, myo-2p::GFP]* | This study |  |
| *C. elegans*: RHS59: *sybIs3970[unc-25p::xbp-1s, myo-2p::GFP]; ldrIs1[dhs-3p::dhs-3::GFP + unc-76(+)]* | This study |  |
| *C. elegans*: RHS60: *sybIs3970[unc-25p::xbp-1s, myo-2p::GFP]; uthSi60[vha-6p::ERss::mRuby::HDEL::unc-54 3'UTR cb-unc-119(+)] IV;* | This study |  |
| *C. elegans*: RHS62: *sybIs3970[unc-25p::xbp-1s, myo-2p::GFP]; pwIs23[vit-2::GFP]* | This study |  |
| *C. elegans*: RHS74: *sybIs3923[eat-4p::xbp-1s, myo-2p::mCherry]; pwIs23[vit-2::GFP]* | This study |  |
| *C. elegans*: RHS79: *sybIs3923[eat-4p::xbp-1s, myo-2p::mCherry]; ldrIs1[dhs-3p::dhs-3::GFP + unc-76(+)]* | This study |  |
| *C. elegans*: RHS84: *sybIs3923[eat-4p::xbp-1s, myo-2p::mCherry]; uthSi60[vha-6p::ERss::mRuby::HDEL::unc-54 3'UTR, cb-unc-119(+)] IV* | This study |  |
| *C. elegans*: RHS96: *sybIs3970[unc-25p::xbp-1s, myo-2p::GFP]; glp-4(bn2) I* | This study |  |
| *C. elegans*: RHS98: *sybIs3923[eat-4p::xbp-1s, myo-2p::mCherry]; glp-4(bn2) I* | This study |  |
| *C. elegans*: RHS109: *uthIs393[vha-6p::Q40::YFP+rol-6(su1006)]; sybIs3923[eat-4p::xbp-1s, myo-2p::mCherry]* | This study |  |
| *C. elegans*: RHS110: *uthIs393[vha-6p::Q40::YFP+rol-6(su1006)]; sybIs3970[unc-25p::xbp-1s, myo-2p::GFP]* | This study |  |
| *C. elegans*: RHS114: *sybIs3954[tbh-1p::xbp-1s, myo-2p::mCherry]; hjSi158[vha-6p::SEL-1(1-79)::mCherry::HDEL::let-858 3’ UTR]* | This study |  |
| *C. elegans*: RHS133: *uthIs393[vha-6p::Q40::YFP+rol-6(su1006)]; sybIs3954[tbh-1p::xbp-1s, myo-2p::mCherry]* | This study |  |
| *C. elegans*: RHS139: *sybIs3954[tbh-1p::xbp-1s, myo-2p::mCherry]; glp-4(bn2) I* | This study |  |
| *C. elegans*: RHS157: *sybIs3970[unc-25p::xbp-1s, myo-2p::GFP]; sqIs61[vha-6p::Q44::YFP + rol-6(su1006)]* | This study |  |
| *C. elegans*: RHS158: *sybIs3923[eat-4p::xbp-1s, myo-2p::mCherry]; sqIs61[vha-6p::Q44::YFP + rol-6(su1006)]* | This study |  |
| *C. elegans*: RHS161: *sybIs3954[tbh-1p::xbp-1s, myo-2p::mCherry]; sqIs61[vha-6p::Q44::YFP + rol-6(su1006)]* | This study |  |
| *C. elegans*: RHS188: *sybIs3924[eat-4p::xbp-1s, myo-2p::mCherry]* | This study |  |
| *C. elegans*: RHS190: *sybIs3924[eat-4p::xbp-1s, myo-2p::mCherry]; uthIs393[vha-6p::Q40::YFP+rol-6(su1006)]* | This study |  |
| *C. elegans*: RHS194: *sybIs3953[tbh-1p::xbp-1s, myo-2p::mCherry]* | This study |  |
| *C. elegans*: RHS196: *sybIs3969[unc-25p::xbp-1s, myo-2p::GFP]* | This study |  |
| *C. elegans*: RHS199: *sybIs3969[unc-25p::xbp-1s, myo-2p::GFP]; uthIs393[vha-6p::Q40::YFP+rol-6(su1006)]* | This study |  |
| *C. elegans*: RHS207: *sybIs3953[tbh-1p::xbp-1s, myo-2p::mCherry]; uthIs393[vha-6p::Q40::YFP+rol-6(su1006)]* | This study |  |
| *C. elegans*: RHS207: *sybIs3953[tbh-1p::xbp-1s, myo-2p::mCherry]; uthIs393[vha-6p::Q40::YFP+rol-6(su1006)]* |  |  |
