## Supplementary material for "Distinct mechanisms of non-autonomous UPR^ER^ mediated by GABAergic, glutamatergic, and octopaminergic neurons": Table S3

| **Primer Name** | **Primer Sequence** | **Primer Purpose** |
| --- | --- | --- |
| xbp-1s RT-qPCR Forward | CGTGCCTTTGAATCAGCAGTG | Measurement of xbp-1s transcripts |
| xbp-1s RT-qPCR Reverse | CGAGGTGTCCATCTTCTTGTT | Measurement of xbp-1s transcripts |
| Y45F10D.4 RT-qPCR Forward | AAGCGTCGGAACAGGAATC | Housekeeping gene |
| Y45F10D.4 RT-qPCR Reverse | TTTTTCCGTTATCGTCGACTC | Housekeeping gene |
| sap-49 RT-qPCR Forward | TGGCGGATCGTCGTGCTTCC | Housekeeping gene |
| sap-49 RT-qPCR Reverse | ACGAGTCTCCTCGTTCGTCCCA | Housekeeping gene |
